# TORC1 and PKA activity towards ribosome biogenesis oscillates in synchrony with the budding yeast cell cycle

**DOI:** 10.1101/2021.05.31.446450

**Authors:** Paolo Guerra, Luc-Alban P.E. Vuillemenot, Marije Been, Andreas Milias-Argeitis

## Abstract

Recent studies have revealed that the growth rate of budding yeast and mammalian cells varies during the cell cycle. By linking a multitude of signals to cell growth, the highly conserved Target of Rapamycin Complex 1 (TORC1) and Protein Kinase A (PKA) pathways are prime candidates for mediating the dynamic coupling between growth and division. However, measurements of TORC1 and PKA activity during the cell cycle are still lacking. Following the localization dynamics of two TORC1 and PKA targets via time-lapse microscopy in hundreds of yeast cells, we found that the activity of these pathways towards ribosome biogenesis fluctuates in synchrony with the cell cycle even under constant external conditions. Mutations of upstream TORC1 and PKA regulators suggested that internal metabolic signals partially mediate these activity changes. Our study reveals a new aspect of TORC1 and PKA signaling, which will be important for understanding growth regulation during the cell cycle.

## Introduction

Cell growth, the collection of processes through which cells accumulate biomass and increase in size, is essential for progression through the cell division cycle (Jorgensen and Tyers, 2004). However, while it has been well-established that the cell cycle involves a complex temporal interplay of many different components, the prevailing notion has been that growth has a much lower temporal complexity, taking place at a constant or exponential rate throughout the cell cycle (Bryan et al., 2012; Elliott et al., 1979a; Elliott and McLaughlin, 1978; Shulman et al., 1973a; Talia et al., 2007). This notion, which is mostly supported by experimental evidence produced decades ago, has been challenged by studies with the model eukaryote *Saccharomyces cerevisiae* (budding yeast). As these works demonstrated, the volume increase rate of budding yeast cells displays a distinct oscillatory pattern during the cell cycle, with maxima during G1 and G2 and minima around the moment of budding and in mitosis (Cookson et al., 2010; Ferrezuelo et al., 2012). Furthermore, budding yeast cell density has been shown to reach a maximum prior to bud formation (Bryan et al., 2010), while autonomous growth rate oscillations coupled to the cell cycle of mammalian cells were also reported recently (Liu et al., 2020). Collectively, these studies have shown that temporal dynamics of cell growth appears to be more complex than was earlier thought.

The target of rapamycin (TOR) and protein kinase A (PKA) pathways are the two main, evolutionarily conserved regulators of cell growth in budding yeast, coupling carbon and nitrogen availability, internal metabolic signals and the presence of noxious stressors to several anabolic and catabolic processes that ultimately impact cell growth and division (Conrad et al., 2014; De Virgilio and Loewith, 2006; Filteau et al., 2015; González and Hall, 2017; Loewith and Hall, 2011; Zurita-Martinez and Cardenas, 2005). Importantly, TOR complex 1 (TORC1) and PKA together regulate ribosome biogenesis and protein synthesis (Jorgensen et al., 2004; Li et al., 2006; Lippman and Broach, 2009; Martin and Hall, 2005; Powers and Walter, 1999; Urban et al., 2007; Yerlikaya et al., 2016), two of the most resource-intensive anabolic processes necessary for biomass accumulation and cell cycle progression (Kafri et al., 2016; Warner, 1999). TORC1 and PKA phosphorylate highly overlapping sets of targets involved in the regulation of ribosomal proteins (RPs), ribosome biogenesis (Ribi) factors and ribosomal RNAs via all three RNA polymerases (Soulard et al., 2010; Wullschleger et al., 2006). By regulating ribosome biogenesis, TORC1 (and likely PKA) also control cytoplasmic crowding and density (Delarue et al., 2018; Neurohr and Amon, 2020). For this reason, these two signaling pathways could be involved in the generation of the non-monotonic growth rate dynamics observed during the cell cycle.

Some connections of TORC1/PKA with the budding yeast cell cycle are known: besides their involvement in ribosome biogenesis, TORC1 and PKA have been implicated in the regulation of the G1/S transition by regulating the abundance of the early G1 cyclin Cln3 (Barbet et al., 1996; Hall, 1998; Mizunuma et al., 2013; Polymenis and Schmidt, 1997). They further antagonize the activity of the Rim15 kinase which controls entry to G0 (Pedruzzi et al., 2003; Reinders et al., 1998; Schneper et al., 2004; Wanke et al., 2005), while TORC1 promotes G1/S transition by stimulating the degradation of the CDK inhibitor Sic1 (Moreno-Torres et al., 2017, 2015; Zinzalla et al., 2007), and is also involved in the regulation of G2/M transition (Nakashima et al., 2008; Tran et al., 2010). Furthermore, polarized growth of yeast cells induced by mating pheromone or apical bud growth in cells deficient in G1/S inhibitor degradation, has been shown to suppress TORC1 activity towards protein synthesis and ribosome biogenesis (Goranov et al., 2013). Beyond yeast, mammalian TORC1 (mTORC1) is also repressed during mitosis (Odle et al., 2020). On the other hand, measurements of key downstream processes such as ribosome biogenesis in yeast have provided contradictory results, with different works proposing that ribosomal protein synthesis proceeds at a constant or exponentially increasing rate during the cell cycle. (Elliott et al., 1979b; Shulman et al., 1973b) Therefore, despite evidence suggesting an interaction of TORC1 and PKA with the cell cycle, we still do not know the activity dynamics of these pathways during the cell cycle, as cell cycle-resolved measurements of TORC1/PKA activity, both in yeast and higher eukaryotes are missing.

Motivated by the observations of growth rate and cell density oscillations, as well as the modulation of growth signaling activity by changes in cell morphology, we sought to investigate if the activities of TORC1 and/or PKA pathways oscillate during the yeast cell cycle. Such a study is challenging to carry out with bulk methods (e.g. Western blots or mass spectrometry) which require synchronous cell populations: invasive synchronization techniques such as arrest-and-release can generate unwanted perturbations in TORC1/PKA activity, while less invasive approaches such as centrifugal elutriation may miss changes in pathway activity due to limited time resolution and/or population synchrony. On the other hand, the lack of reliable single-cell TORC1/PKA activity readouts in yeast has prevented the microscopic study of these pathways during the cell cycle.

To avoid any potential artifacts of population-level methods, we employed single-cell time-lapse fluorescence microscopy of unperturbed dividing cells. This approach does not require synchronization, while offering the possibility to align single-cell traces at different points during the cell cycle, and thus obtain a clear view of cell cycle-regulated processes at high time resolution. Given the central role of ribosomes in biomass accumulation (Lange and Heijnen, 2001), we focused on the ribosome biogenesis (Ribi) branch downstream of TORC1/PKA and studied both pathways together due to the high degree of overlap of TORC1 and PKA targets in the Ribi network. To monitor TORC1/PKA activity via microscopy, we screened transcriptional regulators of RP/Ribi expression whose nuclear localization responds to TORC1/PKA activity, and concluded that the nuclear-to-cytosolic ratio of the Sfp1 activator and Tod6 repressor can serve as a sensitive and fast TORC1 and PKA activity readout. The localization dynamics of Sfp1 and Tod6 during unperturbed growth suggested that TORC1 and PKA activities towards ribosome biogenesis oscillates during the cell cycle, showing a maximum during G1 and a minimum at budding and late mitosis. Analysis of mutants suggested that upstream nutrient-sensing regulators of TORC1 and PKA are part of the mechanism that generates the activity oscillations. Finally, single-cell observations of two fluorescently tagged ribosomal proteins showed that the RP synthesis rate displays strong oscillations which reflect the TORC1/PKA activity pattern. Our findings demonstrate that the activity of these central growth control pathways is temporally organized and tightly coordinated with the cell cycle. This new aspect of TORC1 and PKA signaling will be important for understanding the full scope of cellular activities regulated by these pathways and the mechanisms that couple growth and cell cycle progression.

## Results

### Sfp1 and Tod6 localization is a sensitive and fast reporter of TORC1 and PKA activity

To follow TORC1 and PKA activity in the ribosome biogenesis branch, we searched for TORC1 and PKA targets whose signaling-dependent localization can be monitored via fluorescence time-lapse microscopy. We therefore focused on transcriptional regulators of ribosome biogenesis whose intracellular localization is regulated by TORC1 and/or PKA. A member of this group is the split zinc-finger protein Sfp1 (a functional analog of the c-Myc proto-oncogene (Cook and Tyers, 2007; Lempiäinen et al., 2009)), a direct TORC1 substrate and central activator of hundreds of RP and Ribi genes in response to nutrient and stress (Albert et al., 2018; Reja et al., 2015). Previous studies exploited the fact that the nuclear localization of Sfp1 is regulated by both TORC1 and PKA (Goranov et al., 2013; Jorgensen et al., 2004; Lempiäinen et al., 2009; Marion et al., 2004) and used it as a readout of TORC1/PKA activity (Kingsbury and Cardenas, 2016; Saad et al., 2017; Singh and Tyers, 2009). Phosphorylation of Sfp1 by TORC1 increases the nuclear concentration of the protein and its binding to RP promoters (Albert et al., 2018; Jorgensen et al., 2004; Marion et al., 2004). Much less is known about the regulation of Sfp1 by PKA, besides the fact that active PKA promotes the nuclear accumulation of Sfp1 (Marion et al., 2004; Singh and Tyers, 2009).

Given the lack of a “gold standard” for monitoring the activity of TORC1/PKA via microscopy, we also sought a TORC1/PKA readout complementary to Sfp1, so that one readout could be validated against the other. We therefore screened fluorescent fusions of three repressors of Ribi expression, Tod6, Dot6 and Stb3 (Huber et al., 2011, 2009; Liko, 2007; Liko et al., 2010), as well as the RP repressor Crf1 (Martin et al., 2004). The latter was undetectable via microscopy. Fluorescently tagged Stb3 was visible (Fig.S1), but it was almost fully cytosolic under normal growth, which would compromise the precise quantification of variations in its nuclear-to-cytosolic (N/C) ratio during the cell cycle. Dot6 and Tod6 were both visible and showed an intermediate N/C ratio (Fig.S1). Since Tod6 and Dot6 are paralogs and thus highly similar, we singled out Tod6 for further investigation. Tod6 is a Myb-like helix-turn-helix (HTH) repressor of Ribi expression (Huber et al., 2011, 2009). It is directly phosphorylated and deactivated by the Sch9 kinase (itself a direct target of TORC1 (Urban et al., 2007)) and PKA at multiple Sch9/PKA consensus sites (R[R/K]xS) (Huber et al., 2011, 2009) but, contrary to Sfp1, phosphorylation of Tod6 is thought to promote its nuclear *exit*, though the evidence for this regulation remains circumstantial (Huber et al., 2011, 2009). It is also unclear how Tod6 localization responds to perturbations of TORC1 and PKA pathways. Fig.1A summarizes the current model of Sfp1 and Tod6 regulation by TORC1 and PKA.

**Fig. 1:**
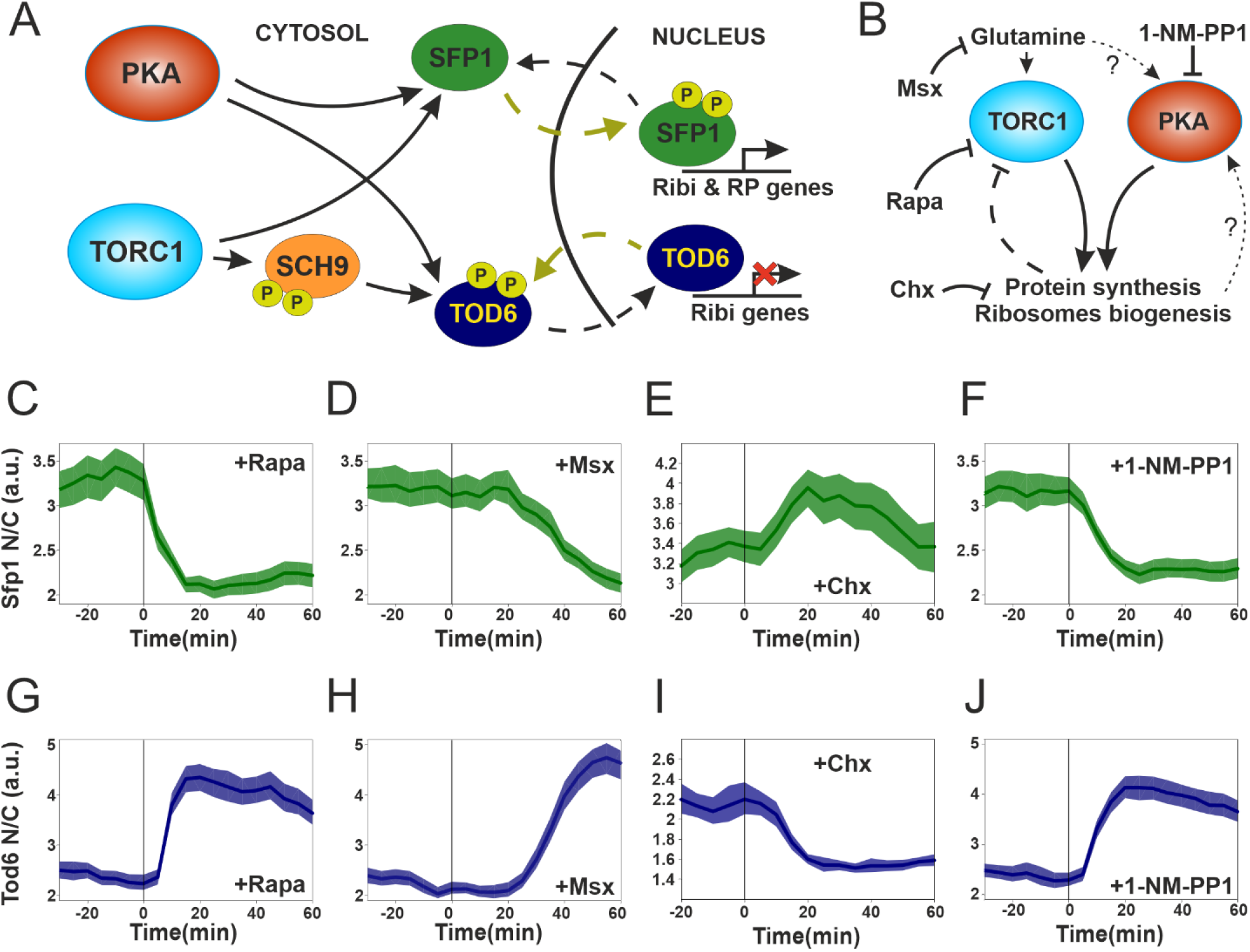
Sfp1 and Tod6 localization is a sensitive and fast reporter of TORC1 and PKA activity. **A**. TORC1 and PKA regulate the phosphorylation and localization of Sfp1 and Tod6. **B**. Schematic representation and summary of TORC1 and PKA chemical perturbations used in this work. **C-F** Sfp1 localization dynamics in response to perturbations shown in **B**. Cells were attached to plastic wells treated with ConcanavalinA (1mg/ml) and imaged every 5 min. Rapamycin (in DMSO) (200ng/ml final), Msx (2mM final), Chx (25µg/ml final) and 1-NM-PP1 (in DMSO) (500nM final) were added at time t = 0 (vertical line). The bands denote the 95% confidence interval for the mean. **G-J** Tod6 localization dynamics in response to perturbations shown in **B**. Cells were attached to plastic wells treated with ConcanavalinA (1mg/ml) and imaged every 5 min. Rapamycin (in DMSO) (200ng/ml final), Msx (2mM final), Chx (25µg/ml final) and 1-

To monitor the nuclear localization of Sfp1 and Tod6 in live cells, we tagged them with a pH-stable tandem GFP [58] and quantified the GFP N/C ratio in cells carrying a fluorescently tagged histone (Hta2-mRFP) as a nuclear marker (Methods and Fig.S2). C-terminal tagging of Sfp1 has been used in several past studies (Goranov et al., 2013; Jorgensen et al., 2004; Kingsbury and Cardenas, 2016; Lempiäinen et al., 2009; Marion et al., 2004; Saad et al., 2017; Singh and Tyers, 2009), and we verified that our tag causes just a minor increase in the cell doubling time, and no changes in cell size (Fig.S1). C-terminal tagging of Tod6 has also been used, though to a lesser extent (Granados et al., 2018; Santos et al., 2019), and we verified that it had no effect on the doubling time or cell size (Fig.S1). These results suggest that the function of both transcription factors is not altered by tagging.

To investigate whether changes in the N/C ratio of Sfp1 and Tod6 can be used as a dynamic readout of TORC1 and PKA activity, we monitored Sfp1 and Tod6 localization dynamics in response to acute perturbations of TORC1 and PKA by rapamycin, methionine sulfoximine, cycloheximide and the bulky ATP-analog 1NM-PP1 (Fig. 1B). Methionine sulfoximine (MSX) is a tightly binding and specific inhibitor of glutamine synthetase, which causes the depletion of intracellular glutamine (a key metabolic input to TORC1 (Stracka et al., 2014; Ukai et al., 2018)), inhibits growth and causes a strong reduction in TORC1 activity towards nitrogen catabolite repression (Crespo et al., 2002; Stracka et al., 2014). Cycloheximide (CHX) blocks translation, while 1-NM-PP1 is a specific inhibitor of kinases carrying a point mutation that enlarges their ATP-binding pocket (Bishop et al., 2000).

In agreement with previous reports (Jorgensen et al., 2004; Marion et al., 2004), Sfp1 moved rapidly out of the nucleus upon inhibition of TORC1 with rapamycin, showing significant delocalization after 10 min (Fig.1C, Fig.S1). Upon MSX treatment, we observed a decrease in the Sfp1 N/C ratio within 20 min, in agreement with the reported timescale for the depletion of the glutamine pool. (Fig.1D) (Crespo et al., 2002). CHX treatment was previously reported to cause a transient increase in TORC1 activity (Jorgensen et al., 2004; Lempiäinen et al., 2009; Santos et al., 2019; Urban et al., 2007) via a still-unexplored negative feedback mechanism (Fig.1B) (Eltschinger and Loewith, 2016). In line with this evidence, we also observed an increase in the Sfp1 N/C ratio 20 min after the addition of CHX (Fig.1E). Even though rapamycin specifically inhibits TORC1, it should be noted that both MSX and CHX cause large metabolic rearrangements which could also alter PKA activity. Thus, to specifically inhibit PKA, we mutated its three kinase subunits (Tpk1,2,3), rendering them sensitive to 1-NM-PP1 (PKAas) (Zaman et al., 2009). 10 min after the addition of 1-NM-PP1 to PKAas cells, we observed a decrease in the N/C ratio of Sfp1 with dynamics and magnitude very similar to the rapamycin response, which demonstrates that PKA is directly involved in the regulation of Sfp1 localization (Fig.1F). These results collectively suggest that Sfp1 localization reports dynamic changes in TORC1 and PKA activity.

Given the fact that active TORC1 and/or PKA are thought to keep Tod6 (a transcriptional repressor) out of the nucleus, we anticipated Tod6 nuclear localization to show an inverse relationship with TORC1 and PKA activity (Fig. 1A,B). In that case, the Tod6 N/C ratio should increase upon inhibition of TORC1 or PKA (i.e. with rapamycin, MSX and 1-NM-PP1) and decrease with hyper-activation of TORC1 by CHX. We indeed observed a marked increase in the Tod6 nuclear localization 10 min after inhibition of TORC1 with rapamycin (Fig.1G, Fig.S1). Treating cells with MSX also caused an increase in the Tod6 N/C ratio with dynamics mirroring the Sfp1 response (Fig.1H). In line with the Sfp1 results and with recently reported observations (Santos et al., 2019), CHX treatment also generated a drop in the Tod6 N/C ratio (Fig.1I), which is consistent with a transient increase in TORC1 activity. Finally, inhibition of PKA by addition of the ATP-competitive inhibitor 1-NM-PP1 to cells carrying the analog-sensitive PKAas mutant caused a fast and strong increase in Tod6 nuclear concentration, very similar to rapamycin treatment (Fig.1J). These results suggest that the nuclear localization of Tod6 is inversely related to TORC1 and PKA activity.

Collectively, our perturbation experiments show that Sfp1 and Tod6 respond sensitively, quickly and on very similar timescales to abrupt changes in TORC1 and PKA activity. They could therefore serve as good proxies of TORC1 and PKA activity during an unperturbed cell cycle. NM-PP1 (in DMSO) (500nM final) were added at time t = 0 (vertical line). The bands denote the 95% confidence interval for the mean.

### Sfp1 and Tod6 localization oscillates in synchrony with the cell cycle

Having established Sfp1 and Tod6 localization as a live-cell readout for TORC1/PKA activity, we sought to investigate how the activity of these pathways may vary during an unperturbed cell cycle by monitoring the localization dynamics of these two proteins. To observe potential fluctuations in Sfp1 and Tod6 localization, we grew cells under constant nutrient conditions (Methods) and imaged them for several hours at high temporal resolution (5 min.). Using Hta2-mRFP as a nuclear marker, we quantified the Sfp1 and Tod6 N/C ratio in single cells over several cell cycles. As cell cycle indicators, we recorded the earliest moment of bud appearance and karyokinesis, indicated by the split of nuclei labelled with Hta2-mRFP (Fig.2A). In the nutrient conditions used, cytokinesis follows karyokinesis after around 5 min (Garmendia-Torres et al., 2018), while the appearance of the bud implies the end of G1. Therefore, the interval between karyokinesis and bud appearance can be used to approximately locate the G1 phase of the cell cycle, while the interval between budding and karyokinesis roughly corresponds to S/G2/M (cf. Methods for further details). We then aligned the individual cell cycle trajectories at karyokinesis and bud appearance, which allowed us to analyze the average localization trends on a common (relative) time axis (Methods).

**Fig. 2:**
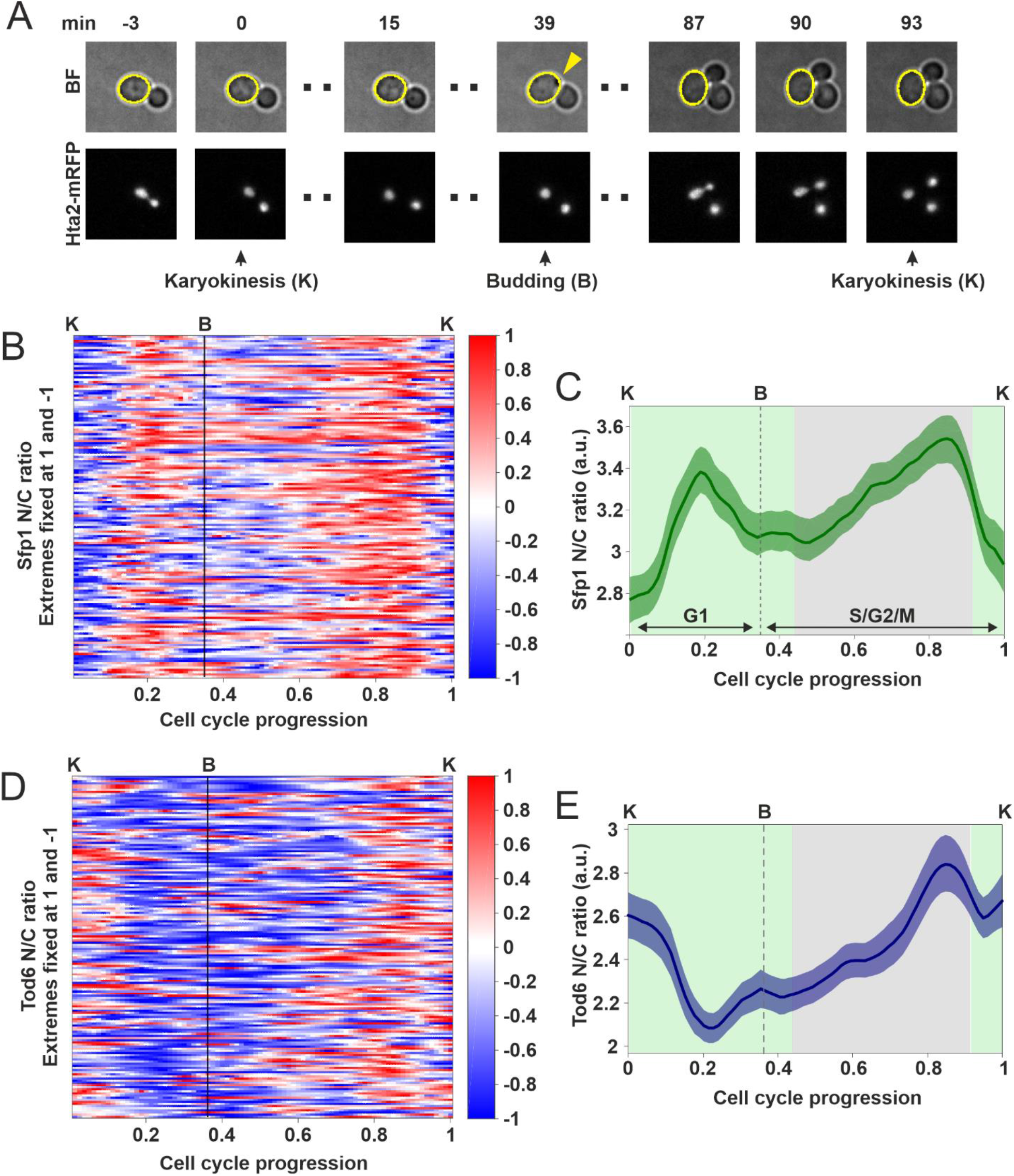
Sfp1 and Tod6 localization oscillates in synchrony with the cell cycle. **A**. Cell cycle events used to manually divide N/C ratio time series into segments corresponding to distinct cell cycles. Karyokinesis (K) was detected with a nuclear fluorescent marker (Hta2-mRFP), and bud appearance (B) corresponded to the earliest moment of a visually detectable membrane deformation in brightfield cell images (indicated by the yellow arrow). Single-cell N/C ratio traces corresponding to individual cell cycles were aligned at three points: a first karyokinesis, bud appearance and the subsequent karyokinesis. **B**. Heatmap of Sfp1 N/C ratio in individual cell cycles (n = 149). Single-cell traces of Sfp1 N/C ratio were aligned and interpolated as described in Fig. 2A and in Methods. For each cell cycle, the Sfp1 N/C ratio was normalized by assigning its maximum to 1 and its minimum to -1, to facilitate the identification of peaks and troughs. **C**. Average Sfp1 N/C ratio dynamics for the cells shown in **B**. The averages were calculated without normalization of the single-cell data. The bands denote the 95% confidence interval for the mean. The average profiles of Sfp1 N/C ratio showed amplitudes in the order of +/-10% around the mean, though single-cell trajectories typically display an even larger amplitude. Sfp1-mNeonGreen showed the same oscillatory pattern (Fig.S3), suggesting that this behavior is not tag-specific **D**. Heatmap of Tod6 N/C ratio in individual cell cycles (n = 161). Single-cell traces of Tod6 N/C ratio were aligned and interpolated as described in Fig. 2A and in Methods. For each cell cycle, the Tod6 N/C ratio was normalized by assigning its maximum to 1 and its minimum to -1, to facilitate the identification of peaks and troughs. **E**. Average Tod6 N/C ratio dynamics for the cells shown in **D**. The averages were calculated without normalization of the single-cell data. The bands denote the 95% confidence interval for the mean. The average profiles of Tod6 N/C ratio showed amplitudes in the order of +/-10% around the mean, though single-cell trajectories typically display an even larger amplitude. Similar to Sfp1, Tod6-mNeonGreen localization dynamics showed the same oscillatory pattern (Fig.S3). In panels **C** and **E** the green background indicates the intervals in which the Sfp1 and Tod6 localization patterns are inverse of each other, whereas the grey background indicates the interval in which Sfp1 and Tod6 localization shows the same behavior.

We observed oscillations in the Sfp1-GFP N/C ratio, both at the single-cell level and the population average (Fig. 2B,C). Sfp1 localization displayed two maxima, one in G1 and one in G2/M, and two minima at budding and karyokinesis (Fig. 2B,C). Tod6-GFP also displayed oscillations in localization with a minimum in G1, an increase trough S/G2/M with a peak in late G2/M and a high N/C ratio through karyokinesis (Fig.2D,E). The inverse patterns of Sfp1 and Tod6 localization during karyokinesis, G1 and early S/G2 (green shaded areas in Fig. 2C, E) are consistent with the inverse responses observed in the chemical perturbations of TORC1 and PKA, showing that the localization of these proteins is likely controlled by TORC1 and PKA in those phases. The observed localization patterns indicate a TORC1/PKA activity peak in G1 and lower activity around karyokinesis and early S/G2. On the other hand, both Sfp1 and Tod6 showed a common increase in average localization during G2/M, which cannot be explained given the inverse relationship between the two readouts and thus needed closer examination.

### Changes in Tod6 localization during G2/M are not caused by TORC1/PKA activity

To investigate if the localization of Tod6 in G2/M is controlled by TORC1 or PKA, we first turned to Sch9, the kinase which relays the signal from TORC1 to Tod6 (Huber et al., 2011), as there is no evidence of a direct TORC1-Tod6 interaction. We replaced the endogenous Sch9 protein by a mutant in which the residues phosphorylated by TORC1 have been substituted to acidic amino acids that mimic constitutive phosphorylation (Sch9_2d3e) (Urban et al., 2007). In this mutant strain, Sch9 regulation is decoupled from TORC1, meaning that inputs from Sch9 to Tod6 are no longer associated with changes in TORC1 activity (Fig.3A). Indeed, in the Sch9_2d3e background Tod6 localization no longer responded to inhibition of TORC1 by rapamycin (Fig.S4), contrary to Sfp1 which still responded normally (Fig.S4).

**Fig. 3:**
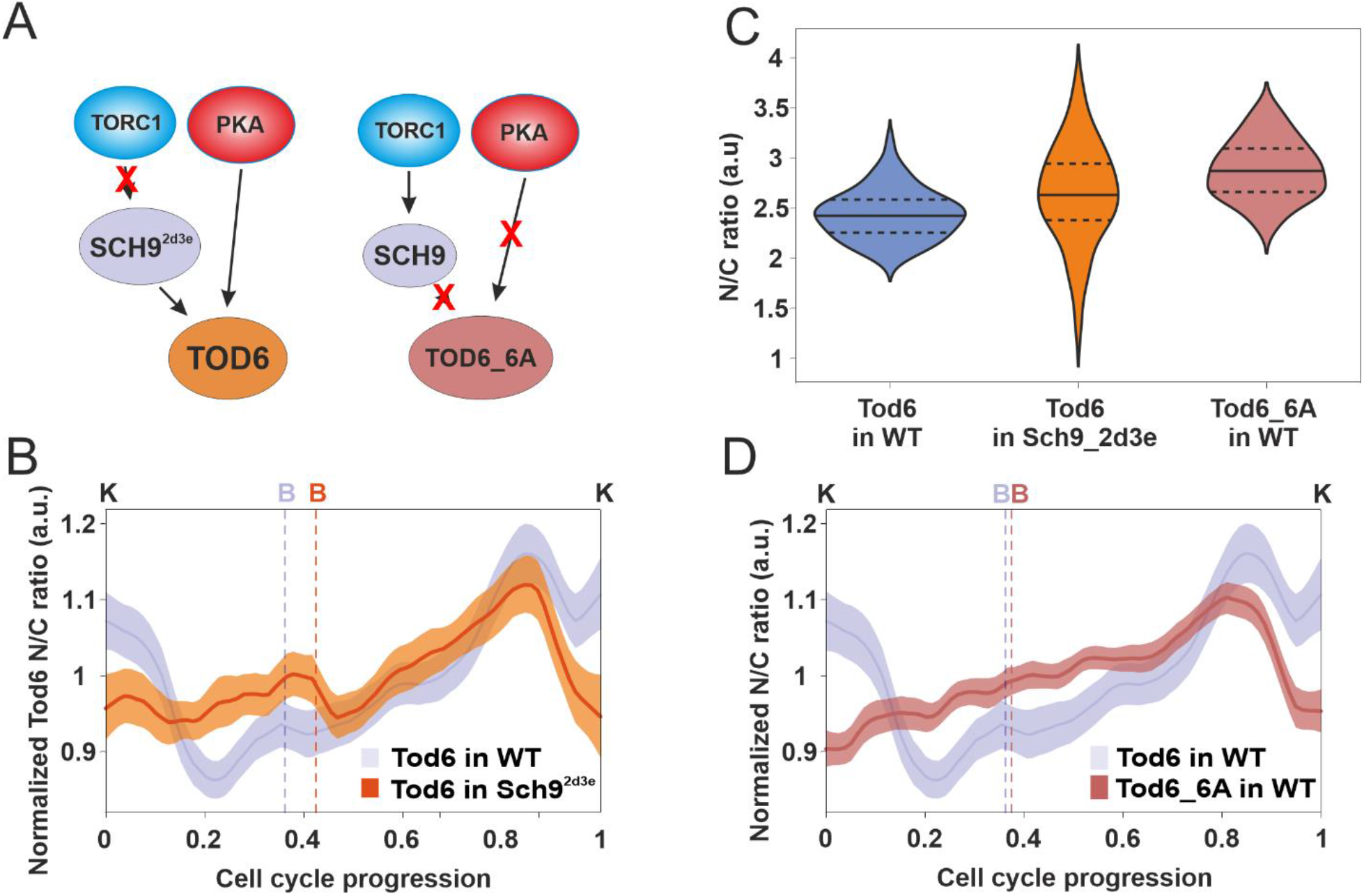
Changes in Tod6 localization during G2/M are not caused by TORC1 or, PKA activity. **A**. Altered regulation of Sch9_2d3e and Tod_6A with respect to wild-type. Left: the phosphomimetic mutations at the Sch9 sites targeted by TORC1 render Sch9 independent from TORC1. Right: alanine substitutions at the six Tod6 sites known to be targeted by Sch9/PKA decouple Tod6 regulation from TORC1 and PKA. **B**. Average Tod6 N/C ratio dynamics in the Sch9_2d3e background (n=103) compared to the wild-type dynamics. The averages were calculated after normalizing each cell cycle trace by its mean, since the average localization of Tod6 differs in the two strains. Cell cycle traces were interpolated and aligned as described in Fig.2A. Bands denote the 95% confidence interval for the mean. **C**. Tod6 N/C ratio distributions in the wild-type (n=70), and the Sch9_2d3e (n=63) and Tod6_6A (n=70) mutants. Median (continuous line) and 25th and 75th percentiles (dashed lines) are also displayed. Statistical comparison, two-tailed Mann-Whitney test: Sch9_2d3e-WT *p* = 8.5·10^−4^, effect size (rank-biserial correlation) *r* = 0.39, Tod6_6A-WT *p* = 2.5·10^−14^, *r* = 0.78. **D**. Average Tod6_6A N/C ratio dynamics (n=109) compared to the wild-type Tod6 dynamics. The averages were calculated after normalizing each cell cycle trace by its mean, since the average localization levels of Tod6 and Tod6_6A differ. Cell cycle traces were interpolated and aligned as described in Fig.2A. Bands denote the 95% confidence interval for the mean.

Quantification of the Tod6 N/C ratio during the cell cycle of the Sch9_2d3e strain showed that all features of the wild-type Tod6 N/C ratio during G1, budding and karyokinesis had disappeared, but the peak in G2/M was still present (Fig. 3B and Fig. S4). This observation indicates that Tod6 localization dynamics during G1, budding and karyokinesis are caused by changes in TORC1 activity, while the localization of Tod6 in G2/M is not controlled by TORC1.

We next asked whether Tod6 localization during G2/M is controlled by PKA or TORC1-independent Sch9 activity. To investigate this, we monitored the localization pattern of a GFP-tagged Tod6 mutant on which all known Sch9 and PKA phosphorylation sites were mutated to alanine to prevent phosphorylation (Tod6_6A), making the regulation of this mutant independent of PKA and TORC1 (Huber et al., 2009) (Fig.3A). Since overexpression of Tod6_6A causes a severe slow-growth phenotype (Huber et al., 2009), we expressed Tod6_6A from an additional chromosomally integrated copy driven by the endogenous Tod6 promoter, leaving the wild-type protein in place. We verified by TORC1 and PKAas inhibition and transient overactivation (via rapamycin, 1-NM-PP1 and CHX respectively) that Tod_6A no longer responds to changes in TORC1 and PKA activity (Fig.S5). We also found that the average N/C ratio of Tod6_6A was higher than in the wild-type (Fig.3C), suggesting that unphosphorylated Tod6 accumulates in the nucleus, as expected.

The N/C profile of Tod6_6A during the cell cycle was very similar to that of the Sch9_2d3e strain, lacking the G1, budding and karyokinesis features of the wild-type and showing the same G2/M peak (Fig.3D and Fig.S5).

Taken together, the observations made with the two mutants indicate that the increase in the N/C ratio of wild-type Tod6 during G2/M is neither generated by TORC1 nor by PKA. On the other hand, the lack of wild-type localization features in G1, budding and karyokinesis in these mutants supports the notion that TORC1 and PKA cause the observed wild-type Tod6 localization changes in those phases.

Collectively, our results suggest that the activity of TORC1 and PKA oscillates in synchrony with the cell cycle. Through the comparison of the two readouts and with the help of mutants we could infer that TORC1/PKA activity peaks during G1, and decreases as cells approach budding and late mitosis, while it is still unclear whether a second activity maximum exists in G2/M, as the Sfp1 data suggest.

### TORC1 and PKA activity oscillations are partially mediated by their nutrient-sensing regulators

If TORC1/PKA activity oscillates during the cell cycle, which mechanisms could be responsible for these oscillations? We reasoned that perturbation of components involved in the generation of TORC1/PKA activity oscillations would manifest itself in altered localization patterns of Sfp1/Tod6. The recent observation of metabolic oscillations (Papagiannakis et al., 2017) which are coupled to the cell cycle and synchronous with the growth rate oscillations prompted us to investigate whether upstream nutrient-sensing regulators of TORC1 and PKA could be implicated in the generation of the observed activity patterns. TORC1 responds to internal metabolic signals generated by carbon and nitrogen sources via two mechanisms: one involving the small GTPases Gtr1/Gtr2 and the vacuolar EGO complex (Binda et al., 2009; Prouteau et al., 2017), and another involving the vacuolar protein Pib2 (Brito et al., 2019; González and Hall, 2017; Ukai et al., 2018; Varlakhanova et al., 2017). Joint deletion of Gtr1/2 and Pib2 is lethal (Ukai et al., 2018), though it is unclear if the two mechanisms act independently of each other and which internal metabolic signals they sense (Ukai et al., 2018; Varlakhanova et al., 2017).

To investigate the potential contribution of Gtr1/2 and Pib2 to the TORC1 activity fluctuations during the cell cycle, we generated ΔGtr1,2 and ΔPib2 strains, and monitored the localization dynamics of Sfp1 and Tod6 during their cell cycle. Although the average N/C ratio of Tod6 was slightly increased in ΔGtr1,2 cells, its localization dynamics was very similar to wild-type, implying that TORC1 activity was not greatly affected in this mutant. (Fig.4A,B)(Fig.S6). On the other hand, deletion of Pib2 caused a marked increase in the average Tod6 N/C ratio, indicating a reduction in TORC1 activity (Fig.4A). Moreover, Pib2 deletion also caused significant changes in the Tod6 localization pattern throughout the cell cycle (Fig. 4C)(Fig.S6) suggesting that Pib2 is implicated in the generation of the Tod6 localization pattern. We did not observe any changes in the average N/C ratio of Sfp1 in both ΔGtr1,2 and ΔPib2 cells, suggesting that these deletions mostly impact the TORC1-Sch9 signaling branch that controls Tod6 (Fig.4D,E,F) (Fig.S6).

**Fig. 4:**
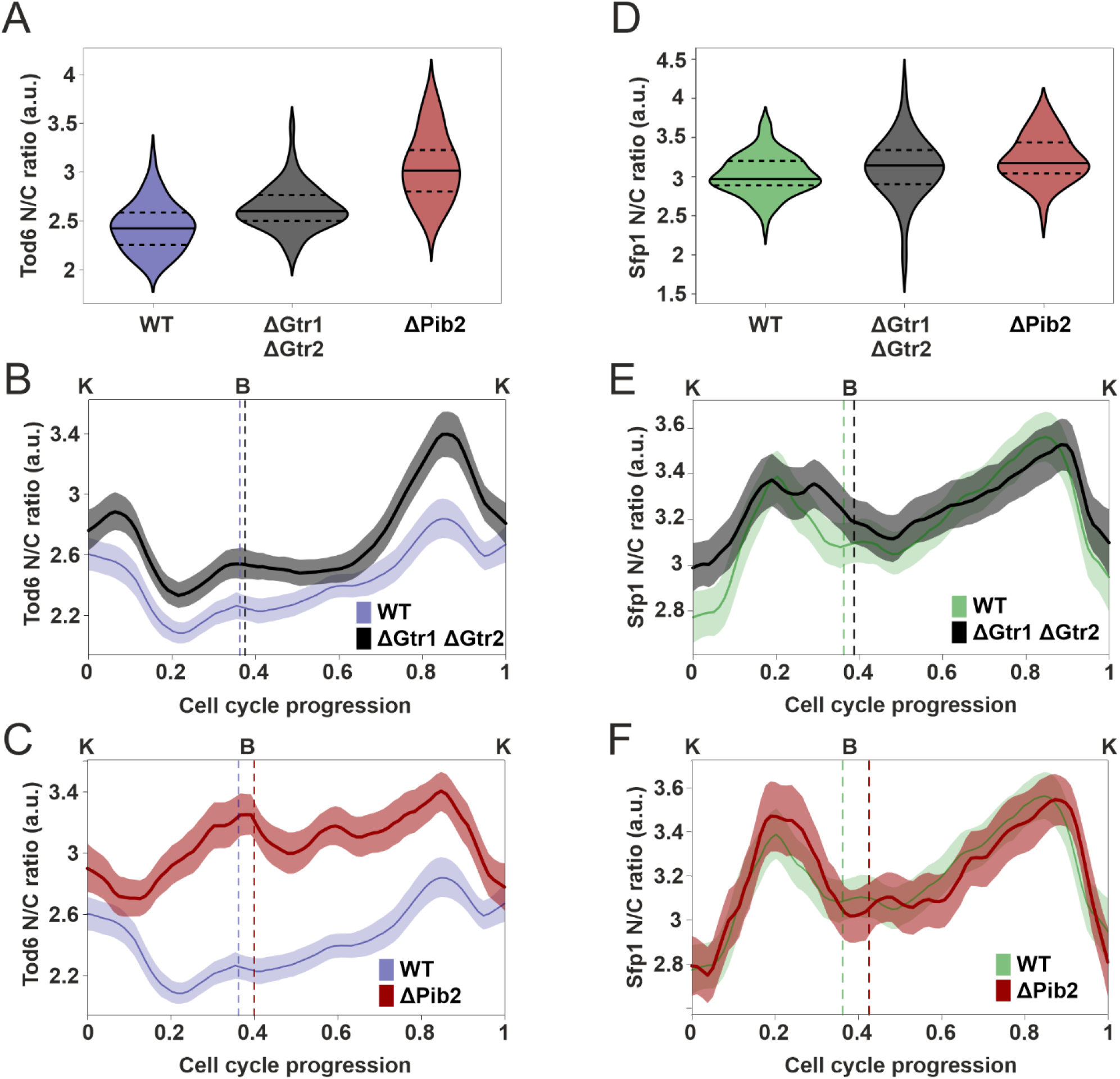
Pib2 mediates TORC1 activity changes towards Tod6 during the cell cycle. **A**. Tod6 N/C ratio distributions in the wild-type (n = 64), and the ΔGtr1ΔGtr2 (n = 57) and ΔPib2 (n = 61) mutants. Median (continuous line) and 25th and 75th percentiles (dashed lines) are also displayed. Statistical comparison, two-tailed Mann-Whitney test: ΔGtr1ΔGtr2-WT *p* = 6.6·10^−6^, r = 0.47, ΔPib2-WT *p* = 6.5·10^−17^, r = 0.86. Effect sizes were larger, especially in ΔPib2 cells. **B**. Average Tod6 N/C ratio dynamics in the ΔGtr1ΔGtr2 background (n = 136) compared to the wild-type (n = 161) dynamics. The averages were calculated without normalization of the single-cell data. Cell cycle traces were interpolated and aligned as described in Fig.2A. Bands denote the 95% confidence interval for the mean. It is interesting to observe that the stronger G2 peak in the N/C dynamics of the mutant strain is a consequence of better synchronization in the single-cell G2 localization pulses of Tod6 (Fig.S6). **C**. Average Tod6 N/C ratio dynamics in the ΔPib2 background (n = 113) compared to the wild-type (n = 161) dynamics. The averages were calculated without normalization of the single-cell data. Cell cycle traces were interpolated and aligned as described in Fig.2A. Bands denote the 95% confidence interval for the mean. **D**. Sfp1 N/C ratio distributions in the wild-type (n = 70), and the ΔGtr1ΔGtr2 (n = 60) and ΔPib2 (n = 43) mutants. Median (continuous line) and 25th and 75th percentiles (dashed lines) are also displayed. Statistical comparison, two-tailed Mann-Whitney test: ΔGtr1ΔGtr2-WT *p* = 0.033, r = 0.21, ΔPib2-WT *p* = 3.5·10^−3^, r = 0.4. Even though the second p-value is quite small, the observed effect (difference in medians) is not large. **E**. Average Sfp1 N/C ratio dynamics in the ΔGtr1ΔGtr2 background (n = 122) compared to the wild-type (n = 149) dynamics. The averages were calculated without normalization of the single-cell data. Cell cycle traces were interpolated and aligned as described in Fig.2A. Bands denote the 95% confidence interval for the mean. **F**. Average Sfp1 N/C ratio dynamics in the ΔPib2 background (n = 79) compared to the wild-type (n = 149) dynamics. The averages were calculated without normalization of the single-cell data. Cell cycle traces were interpolated and aligned as described in Fig.2A. Bands denote the 95% confidence interval for the mean.

To investigate whether Gtr1,2 can compensate for the absence of Pib2, we mutated Gtr1 and Gtr2 to compensate for the apparently deficient TORC1 activation of these cells by inserting point mutations (Gtr1_Q65L and Gtr2_S23L) which lock the Gtr proteins into their TORC1-activating configuration (Gtr1-GTP and Gtr2-GDP) (Binda et al., 2009; Nakashima et al., 1999). However, the Tod6 localization pattern in ΔPib2_Gtr1_Q65L_Gtr2_S23L cells remained very similar to that of ΔPib2 background (Fig.S6), while the average Sfp1 N/C ratio again remained unaffected (Fig.S6).

Altogether, mutations of upstream TORC1 regulators demonstrated that Pib2 is implicated in the generation of the TORC1 activity fluctuations during the cell cycle, whereas this cannot be said for Gtr1,2. It is still possible that Gtr1,2 participate in the pattern generation but their contribution is not visible under the high glucose conditions tested here. Alternatively, the loss of these proteins could somehow be compensated via feedback regulation, which abounds in the TORC1 network (Eltschinger and Loewith, 2016). In any case, the different responses of our readouts upon Pib2 and Gtr1,2 deletions suggests that these two upstream regulators of TORC1 are not entirely interchangeable during unperturbed growth.

Next, we evaluated the effect of upstream PKA regulators. Recently, it was shown that the membrane-bound Ras2 GTPase couples glycolytic flux to PKA activation (Peeters et al., 2017a), while both TORC1 and PKA pathways are implicated in intracellular pH homeostasis via Gtr1/2 and the vacuolar ATPase (Deprez et al., 2021; Oliveira et al., 2015). PKA activity is regulated at several points along the pathway: the activity of the kinase subunits Tpk1,2,3 is repressed by the binding of the regulatory subunit Bcy1, and this interaction is abrogated by the binding of cAMP to Bcy1 (Santangelo, 2006; Toda et al., 1987). The levels of cAMP are in turn controlled by Ras2 and the Gpa2 Gα protein, both of which stimulate the adenylate cyclase Cyr1 (Broach, 1991; Santangelo, 2006).

To determine the contribution of upstream PKA regulators to the observed activity fluctuations, we used hyperactive Tpk kinase mutants: ΔBcy1 and Ras2_A18V19_Gpa2_A273. The first mutant lacks the negative regulator Bcy1 (and thus, most of the known regulation on Tpk1-3) resulting in (constitutively) hyperactive Tpk1,2,3, while the second mutant contained the constitutively active forms of the two known upstream regulators, Ras2 and Gpa2. These mutations also hyperactivate the PKA pathway (Jorgensen et al., 2004; Kraakman et al., 1999; Schmelzle et al., 2004). In both strains, we did not observe a major change in the average N/C ratio of Sfp1, but the amplitude of the Sfp1 N/C oscillations was reduced, especially in ΔBcy1 (Fig.5A,B,C)(Fig.S7). In particular, the minima around budding and karyokinesis (corresponding to low-activity periods of TORC1/PKA in wild-type) were much shallower, in agreement with the fact that PKA is constitutively (hyper)active in these strains throughout the cell cycle. On the other hand, the nuclear localization of Tod6 showed a dramatic decrease, while the N/C oscillations were greatly suppressed and displayed an altered pattern (Fig.5D,E,F)(Fig.S7). These observations are again consistent with a hyperactive PKA throughout the cell cycle.

**Fig. 5:**
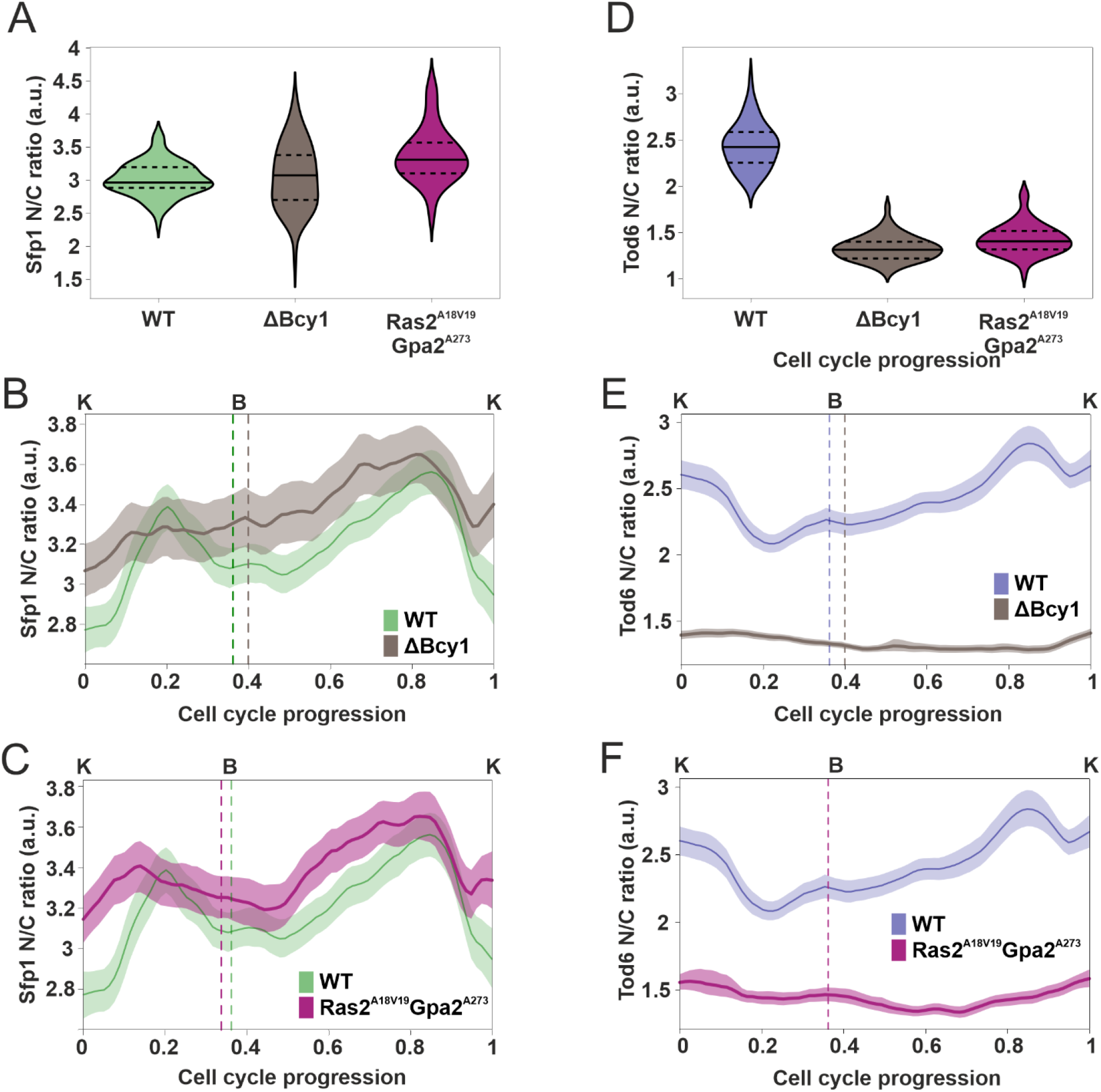
Bcy1 and Ras2/Gpa2 mediate PKA activity changes towards Sfp1 and Tod6 during the cell cycle. **A**. Sfp1 N/C ratio distributions in the wild-type (n = 70) and the two mutants with hyperactive PKA (Bcy1KO: n = 62, Ras2A18V19_Gpa2A273: n = 69). Median (continuous line) and 25th and 75th percentiles (dashed lines) are also displayed. Statistical comparison, two-tailed Mann-Whitney test: ΔBcy1-WT *p* = 0.77, r = 0.03, Ras2_A18V19 Gpa2_A273-WT *p* = 2.1·10^−9^, r = 0.59. **B**. Average Sfp1 N/C ratio dynamics in the ΔBcy1 background (n = 125) compared to the wild-type (n = 149) dynamics. The averages were calculated without normalization of the single-cell data. Cell cycles were interpolated and aligned as described in Fig.2A. Bands denote the 95% confidence interval for the mean. **C**. Average Sfp1 N/C ratio dynamics in the Ras2_A18V19 Gpa2_A273 background (n = 111) (in which Ras2 and Gpa2 are locked in their active forms) compared to the wild-type (n = 149) dynamics. The averages were calculated without normalization of the single-cell data. Cell cycles were interpolated and aligned as described in Fig.2A. Bands denote the 95% confidence interval for the mean. **D**. Tod6 N/C ratio distributions in the wild-type (n = 64) and the two mutants with hyperactive PKA (Bcy1KO: n = 59, Ras2A18V19_Gpa2A273: n = 46). Median (continuous line) and 25th and 75th percentiles (dashed lines) are also displayed. Statistical comparison, two-tailed Mann-Whitney test: ΔBcy1-WT *p* = 1.2·10^−21^, r = 1, Ras2_A18V19 Gpa2_A273-WT *p* = 1.4·10^−17^, r = 0.95. **E**. Average Tod6 N/C ratio dynamics in the ΔBcy1 background (n = 131) compared to the wild-type (n = 161) dynamics. The averages were calculated without normalization of the single-cell data. Cell cycles were interpolated and aligned as described in Fig.2A. Bands denote the 95% confidence interval for the mean. **F**. Average Tod6 N/C ratio dynamics in the Ras2_A18V19 Gpa2_A273 background (n = 100) compared to the wild-type (n = 161) dynamics. The averages were calculated without normalization of the single-cell data. Cell cycles were interpolated and aligned as described in Fig.2A. Bands denote the 95% confidence interval for the mean.

In summary, the perturbations of the main upstream regulators of TORC1 and PKA have generated changes in the cell cycle patterns of our readouts, consistent with the notion that the upstream, nutrient-sensing regulators (Pib2 in the case of TORC1 and Bcy1/Ras2/Gpa2 in the case of PKA) are part of the mechanism that generates the activity fluctuations in TORC1 and PKA during an unperturbed cell cycle.

### Alterations in the TORC1 and PKA cell cycle activity pattern perturb the coupling between cell cycle and growth

Given that mutations upstream of TORC1 and PKA resulted in altered activity dynamics during the cell cycle, we asked whether these altered dynamics are connected with more “global” changes in cell size and growth. To this end, we determined by microscopy the volume distribution of a cell population at two critical cell cycle points (budding and karyokinesis), the volume increase rate during the cell cycle, and lengths of cell cycle phases durations in cells carrying the Pib2 or Bcy1 deletions.

Consistent with the lowered TORC1 activity which we had inferred in ΔPib2 cells (Fig.4A,C), we observed an overall lower volume increase rate in these cells compared to the wild-type (Fig.6A). At the same time, the duration of both growth phases was prolonged (resulting in a small increase in total cell cycle duration) (Fig.6B,C), while mother cells were smaller at the moment of bud appearance, and grew smaller buds than the wild-type (Fig.6E,F). As expected from the Sfp1 and Tod6 N/C ratio profiles (Fig.S6), mutation of Gtr1,2 to their constitutively active form did not fully revert these changes, as it restored the WT cell cycle phase durations, but not the cell volumes (Fig. S8).

**Fig. 6:**
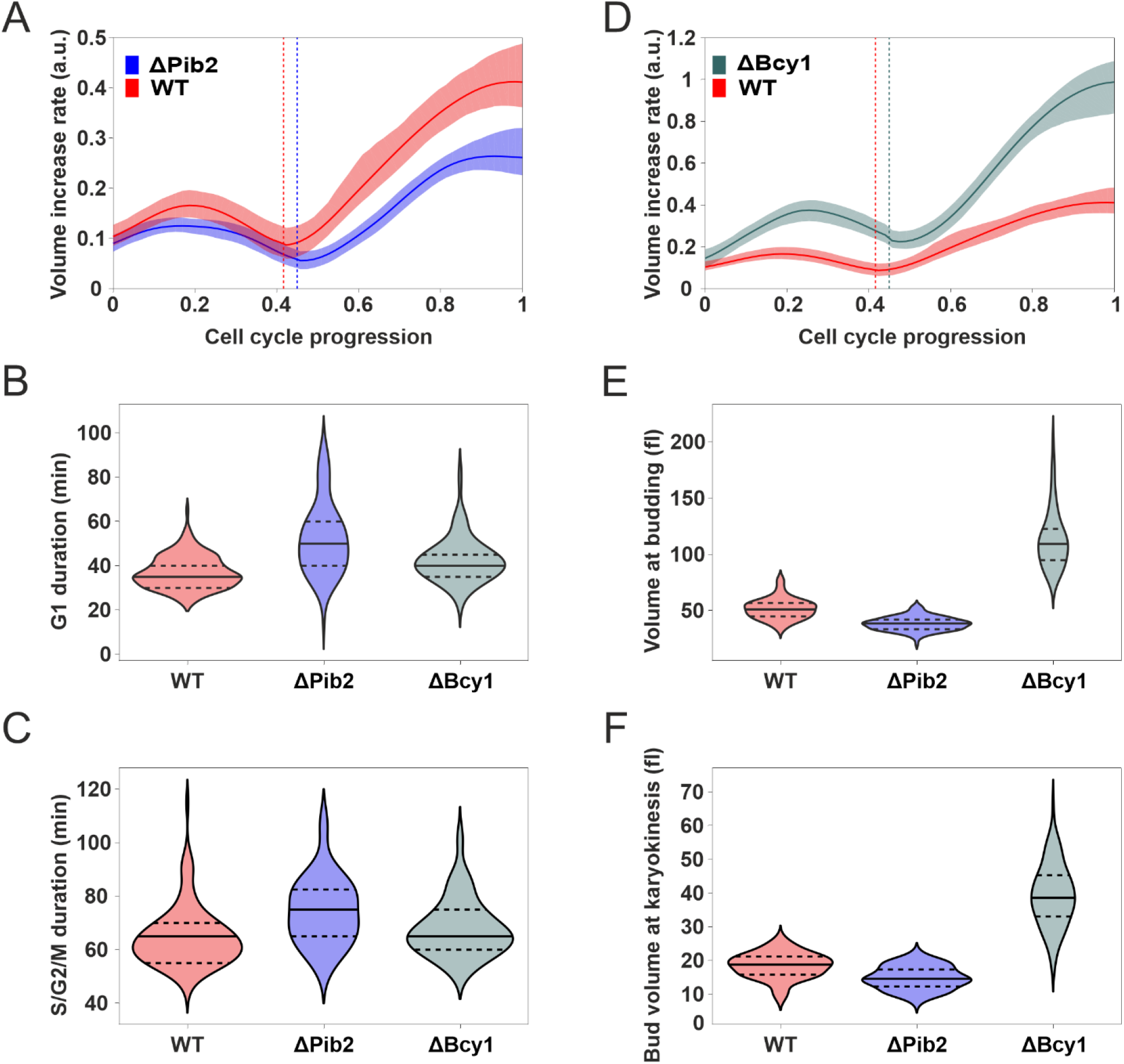
Alterations in the TORC1 and PKA cell cycle activity pattern perturb the coupling between cell cycle and growth. **A**. Average volume increase rate during the cell cycle of the wild-type and ΔPib2 mutant. Volumes of single cells were tracked, manually split into individual cell cycles, smoothed and differentiated (Methods). Data from individual cell cycles traces were interpolated and aligned as described in Fig.2A. Bands denote the 95% confidence interval for the mean. **B**. G1 duration distributions in the wild-type (n = 145) and the ΔPib2 (n = 112) and ΔBcy1 (n = 135) mutants. Median (continuous line) and 25th and 75th percentiles (dashed lines) are also displayed. G1 was defined as the interval between karyokinesis and bud appearance. This definition slightly overestimates the actual G1 duration. Statistical comparison, two-tailed Mann-Whitney test: ΔPib2-WT *p* = 1.5·10^−17^,r = 0.61, ΔBcy1-WT *p* = 4.7·10^−7^, r = 0.34. **C**. S/G2/M duration distributions in the wild-type (n = 113) and the ΔPib2 (n = 103) and ΔBcy1 (n = 113) mutants. Median (continuous line) and 25th and 75th percentiles (dashed lines) are also displayed. S/G2/M was defined as the interval between bud appearance and karyokinesis. This definition slightly underestimates the actual duration of these phases. Statistical comparison, two-tailed Mann-Whitney test: ΔPib2-WT *p* = 1·10^−9^, r = 0.47, ΔBcy1-WT *p* = 0.004, r = 0.22 **D**. Average volume increase rate during the cell cycle of the wild-type and ΔBcy1 mutant. The processing of volume data was carried out as described in **A. E**. Distributions of volume at the moment of bud appearance in the wild-type (n = 150) and the ΔPib2 (n = 122) and ΔBcy1 (n = 135) mutants. Median (continuous line) and 25th and 75th percentiles (dashed lines) are also displayed. Statistical comparison, two-tailed Mann-Whitney test: ΔPib2-WT *p* = 5.6·10^−29^, r = 0.78 ΔBcy1-WT *p* = 5.9·10^−48^, r = 0.99. **F**. Distributions of bud volume at the moment of karyokinesis in the wild-type (n = 109) and the ΔPib2 (n = 103) and ΔBcy1 (n = 110) mutants. Median (continuous line) and 25th and 75th percentiles (dashed lines) are also displayed. Statistical comparison, two-tailed Mann-Whitney test: ΔPib2-WT *p* = 3.2·10^−11^, r = 0.53, ΔBcy1-WT *p* = 1.9·10^−35^, r = 0.97

On the other hand, deletion of Bcy1 resulted in a volume increase rate which was considerably larger than the wild-type (Fig.6D). The difference was evident throughout the cell cycle, and more pronounced in the second half. At same time, the duration of G1 and S/G2/M did not change (Fig.6B,C) and consequently, cells were much larger, budding at higher volumes and growing larger buds than the wild-type (Fig.6E,F). Interestingly, the volume of ΔBcy1 cells displayed much more variability compared to wild-type, which suggested that size homeostasis mechanisms are less effective in these cells, presumably due to the decoupling of PKA regulation from upstream cues. Similar, but slightly less pronounced increases with respect to wild-type were observed in the Ras2-Gpa2 mutant strain (Fig. S8).

Overall, the observations of cell growth in the two mutant strains, demonstrate that, besides altering the TORC1 and PKA activity pattern during the cell cycle, perturbations of nutrient-sensing regulators upstream of TORC1 and PKA also affect the coupling of cell cycle progression and cell size control mechanisms. It is interesting to note that, whereas TORC1 and PKA activity had so far been considered necessary for G1 progression, our results show a clear contribution of these pathways to growth during G2/M.

### The synthesis rate of ribosomal proteins Rpl13a and Rpl26a reflects TORC1 and PKA activity changes during the cell cycle

Having observed that TORC1 and PKA activity towards readouts connected with ribosome biogenesis oscillates in synchrony with the cell cycle, we investigated whether this activity pattern is also reflected in ribosome biogenesis dynamics. To explore how ribosome biogenesis proceeds during the cell cycle, we used the synthesis of ribosomal proteins (RPs) as a proxy. Previous attempts to quantify yeast RP synthesis based on bulk measurements of cultures synchronized via centrifugal elutriation. However, the limitations of bulk synchronization and analysis methods are particularly relevant when studying the cell cycle dynamics of mother cells growing in rich nutrients, where cell cycle duration is the shortest. (Elliott et al., 1979b; Shulman et al., 1973b).

To overcome these limitations, we turned again to time-lapse fluorescence microscopy. To measure RP synthesis dynamics in growing single cells, we tagged with sfGFP two ribosomal proteins, Rpl13a and Rpl26a (tagging of these RPs did not affect cell growth (Fig. S9)). TORC1 has been reported to control Rpl13a expression (Marion et al., 2004), while Rpl26a promoter activity has been used as a TORC1 reporter in liquid cell cultures (Kessi-Pérez et al., 2018).

We measured the total abundance of Rpl13a-GFP and Rpl26a-GFP in single cells using the same cell cycle indicators and alignment procedure as above. For each cell cycle, we recorded the average GFP fluorescence intensity and (mother+bud) volume time traces to obtain total GFP abundance, smoothed the resulting abundance time series, and finally calculated the GFP synthesis rate at each point in time via differentiation and correction for GFP maturation (Methods). Using budding and karyokinesis, we then aligned the single-cell traces from cycles of different length on a common (relative) time axis denoting cell cycle progression (Methods and Fig. 2A).

We found that the synthesis rate of Rpl13a-GFP and Rpl26a-GFP per unit of volume is not constant or exponentially increasing, but presents two maxima (in G1 and in G2/M) and two minima (after budding and during mitosis) (Fig.7A,B). The magnitude of these changes indicates that the synthesis rate of Rpl13a and Rpl26a is strongly regulated during the cell cycle (Fig.7C,D).

**Fig. 7:**
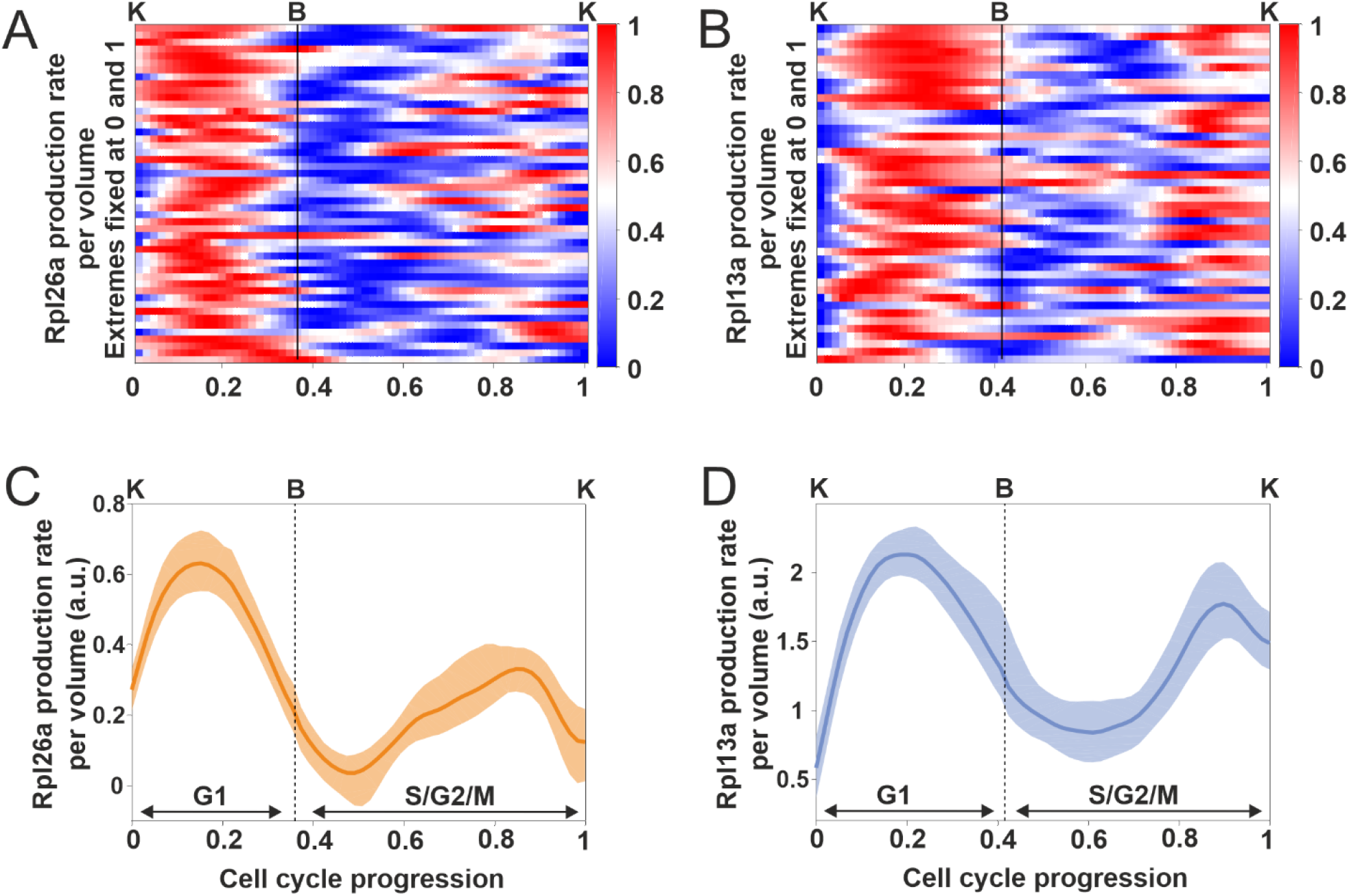
The synthesis rate of the ribosomal proteins Rpl13a and Rpl2a oscillates in synchrony with the cell cycle. **A, B**. Heatmaps of estimates of Rpl26a and Rpl13a synthesis rate per unit of volume in individual cell cycles (n = 44 cell cycles for Rpl13a, n = 49 cell cycles for Rpl26a). Time series of individual cell cycles were interpolated and aligned as described in Fig.2A. For each cell cycle, the time series of the RP synthesis rate was normalized by assigning its maximum to 1 and its minimum to 0, to facilitate the identification of peaks and troughs. Cells were grown in glucose minimal medium over several cell cycles and imaged with a fast frame rate (5min). **C, D** Average Rpl26a and Rpl13a synthesis rate per unit volume for the cells shown in **A** and **B**. The averages were calculated without normalization of the single-cell data. The bands denote the 95% confidence interval for the mean.

The observed RP synthesis dynamics are in good agreement with the inferred TORC1/PKA activity pattern, with the exception of the G2/M peak, which was only suggested by the Sfp1 dynamics and could not be confirmed with Tod6. Given that the TORC1 and PKA pathways are the main regulators of ribosome synthesis (Jorgensen et al., 2004; Lippman and Broach, 2009; Powers and Walter, 1999; Zurita-Martinez and Cardenas, 2005), the determined RP synthesis rate further supports our finding that the activity of these signaling pathways oscillates during the cell cycle.

## Discussion

By using time-lapse fluorescence microscopy and meticulous processing of single-cell data we found evidence that the activity of TORC1 and PKA towards ribosome biogenesis oscillates in synchrony with the cell cycle. Using the intracellular localization of Sfp1 and Tod6 as a proxy for TORC1/PKA activity, we observed coordinated changes in our two readouts in unperturbed cell cycles of individual cells, and found that TORC1/PKA activity shows a maximum during G1 and two minima: one around budding and another in late M/early G1. Meanwhile, it remains unclear what the pattern of TORC1 and PKA activity is during G2. Nutrient-sensing mechanisms upstream of TORC1 and PKA are involved in the regulation of TORC1 and PKA during the cell cycle, suggesting that these signaling pathways respond in part to oscillatory internal metabolic signals during the cell cycle (Amponsah et al., 2021; Baumgartner et al., 2018; Papagiannakis et al., 2017). Perturbations of the TORC1/PKA activity oscillations caused changes in the coupling between growth and cell cycle progression. Finally, the synthesis rate of two ribosomal proteins displayed oscillations in good agreement with the inferred TORC1/PKA oscillations, further supporting the notion that TORC1 and PKA activity oscillates during the cell cycle.

Through the characterization of Sfp1 and Tod6 localization dynamics via chemical perturbation experiments, we obtained new insights into the regulation of these proteins and opened up directions for future work. For example, Sfp1 localization seems to be directly regulated by PKA, given the fast response of Sfp1 localization to the 1NM-PP1 perturbation. This observation is further supported by the fact that an Sfp1 mutant with all known TORC1 sites mutated to alanine (Sfp1-1 (Lempiäinen et al., 2009)) displayed reduced nuclear localization relative to wild-type, but still responded to our chemical perturbations (Fig.S5), confirming earlier observations that this mutant still retains some wild-type functionality, and suggesting that additional functional phosphorylation sites exist on the protein (Lempiäinen et al., 2009) or that its localization is partially regulated through the binding of other proteins (e.g. Mrs6), which themselves are targets of signaling (Lempiäinen et al., 2009; Singh and Tyers, 2009). Besides Sfp1, we demonstrated a connection between Tod6 phosphorylation and localization, showing that the latter responds to changes in TORC1 and PKA activity. Knowing the mechanisms by which TORC1 and PKA individually regulate Sfp1 and Tod6 localization will be instrumental for the development of biosensors specific for TORC1 or PKA in the future.

The inferred peak in TORC1 and PKA activity during G1 is in agreement with their role in regulating G1 progression and the G1/S transition (Barbet et al., 1996; Hall, 1998; Mizunuma et al., 2013; Polymenis and Schmidt, 1997). In particular, we recently showed that transition through the G1 checkpoint (also known as “Start”) is triggered by a pulse in protein synthesis during G1 which causes a pulse in the concentration of the unstable G1 cyclin Cln3 (Litsios et al., 2019). Both TORC1 and PKA have been reported to promote traversal through START by adjusting global protein synthesis (and thus, Cln3 levels) in response to nutrients (Barbet et al., 1996; Hall, 1998; Mizunuma et al., 2013; Polymenis and Schmidt, 1997). Our results may therefore explain the cause of the observed pulses in protein synthesis and Cln3 concentration during G1. The decrease in TORC1/PKA activity around the moment of budding also corroborates previous observations that polarized growth (induced by mating pheromone treatment or in cell cycle mutants) decreases TORC1 activity (Goranov et al., 2013), though it was unknown whether similar downregulation took place during normal cell cycle progression. While we could not resolve TORC1 and PKA activity dynamics in G2/M, the observations of the mutant strains (Fig. 6), as well as the second pulse in RP expression in G2/M indicate that these pathways are active also during G2. Finally, although the budding yeast and mammalian mitosis have several differences, it is interesting to note that the decrease in TORC1/PKA activity around karyokinesis strongly resembles the growth dynamics observed in mammalian cells, where mass accumulation and protein synthesis persist through prophase, slow down as cells approach the metaphase-to-anaphase transition, and recover during late cytokinesis (Miettinen et al., 2019).

Our investigation of TORC1 and PKA activity patterns in response to mutations and deletions in their nutrient-sensing regulators showed that Pib2 deletion causes a significant change in Tod6 localization and has a significant effect on volume dynamics during the cell cycle. The fact that the Sfp1 localization pattern was not altered significantly in this mutant suggests that Pib2 deletion has a larger effect on the Sch9 output branch of TORC1 signaling compared to Sfp1 phosphorylation. The presumed decrease in Tod6 phosphorylation in ΔPib2 cells (demonstrated by an increase in the average Tod6 N/C ratio) would be consistent with the fact that Sch9 phosphorylation (and, hence, activity) shows a clear decrease in ΔPib2 cells compared to wild-type (Ukai et al., 2018). Although both Pib2 and Gtr1/2 were known to modulate TORC1 activity in response to nitrogen source shifts and noxious stressors, it was not clear if they played a role in normal cell cycle progression. Our results suggest that Pib2, a recently discovered sensor of intracellular glutamine in budding yeast, could relay internal metabolic signals that also modulate TORC1 activity during the cell cycle. Upstream regulators of PKA such as Bcy1 and Ras2/Gpa2 are also clearly involved in the modulation of PKA activity, as evidenced by the changes induced on the localization of both Sfp1 and Tod6 upon Bcy1 deletion and the locking of Ras2 and Gpa2. Notably, Bcy1 binding to Tpk1-3 is regulated by cyclic AMP, a central signaling metabolite, while further upstream, Ras2 activity has been shown to respond to changes in glycolytic flux (Peeters et al., 2017b). It remains to be seen precisely which metabolic inputs are in fact responsible for the observed TORC1 and PKA activity changes during each cell cycle phase.

Overall, the results presented in this work demonstrate that the activity of two central pathways controlling cell growth is temporally organized and tightly connected with cell cycle progression. Our observation that TORC1/PKA activity fluctuates in synchrony with the cell cycle is in line with a recent report that metabolic precursors are synthesized at different rates during the cell cycle (e.g. amino acid synthesis increases in G1 and G2/M with a minimum in S) (Campbell et al., 2020), as well as the observation that metabolism operates as an autonomous oscillator that is coupled to the cell cycle (Papagiannakis et al., 2017). On the other hand, it becomes increasingly apparent that growth-controlling signaling pathways are in direct, bidirectional communication with the cell-cycle machinery (Moreno-Torres et al., 2017, 2015; Odle et al., 2020; Romero-Pozuelo et al., 2020; Talarek et al., 2017). It will therefore be instructive to investigate further whether the activity of TORC1 and PKA towards catabolic processes such as autophagy is also coordinated with the cell cycle, as recent evidence suggests (Li et al., 2020; Odle et al., 2020; Yamasaki et al., 2020). Untangling the complex interactions of these fundamental intracellular processes will provide a much deeper understanding of how proliferating cells coordinate growth with division.

## Methods

### Yeast strains

All strains were constructed on the S288C-derived prototrophic YSBN6 background (81). Integration of fluorescent reporters was carried out using homologous recombination. Fragments containing the fluorescent protein, the resistance cassette and the appropriate flanking sequences were amplified from plasmids built via Gibson Assembly. Substitutions of Bcy1 and Pib2 with the resistance cassette were constructed the same way. The sequence for the mutant Tod6_6A(S280A S298A S308A S318A S333A S346A) was taken from plasmid pAH268 (Huber et al., 2009). The sequence for the mutant Sfp1-1(S39A S170A S181A S183A T227A T228A T446A) was taken from plasmid pHL150 (Lempiäinen et al., 2009). The sequence for the mutant Sch9_2D3E (T723D, S726D, T737E, S758E, S765E) was taken from plasmid pJU841 (Urban et al., 2007). Strains containing deletion of Gtr1 or Gtr2 and mutation of Tpk1(M164G), Tpk2(M147G), Tpk3(M165G), Gtr1(Q65L), Gtr2(S23L), Ras2(A18_V19) or Gpa2(A273) were constructed via CRISPR-Cas9 using the MoClo Yeast Toolkit (Lee et al., 2015). Target sequences used for CRISPR-Cas9 mutagenesis and deletions can be found Tabsle S1. All strains were verified by PCR and sequencing.

### Yeast growth media and cultivation

Cells were grown in minimal medium (Verduyn et al., 1992) supplemented with 2% glucose (Sigma-Aldrich). Batch culture was carried out at 30° C with shaking at 300 rpm. Exponentially growing cells were used for microscopy experiments.

### Microscopy

All microscopy experiments were performed using inverted fluorescence microscopes (Eclipse Ti-E, Nikon Instruments). Temperature was kept constant at 30°C using a microscope incubator (Life Imaging Services). For all the experiments a 100x Nikon S Fluor (N.A.=1.30) objective was used. Images were recorded using iXon Ultra 897 DU-897-U-CD0-#EX cameras (Andor Technology). Fluorescence measurements were performed using an LED-based excitation system (pE2; CoolLED Limited and Lumencor, AURA). For GFP measurements cells were excited at 470nm (Excitation filter: 450-490nm, dichroic: 495nm, emission filter: 500-550nm). For YFP measurements cells were excited at 500nm (Excitation filter: 490-510nm, dichroic: 515nm, emission filter: 520-550nm). For RFP measurements cells were excited at 565nm (Excitation filter: 540-580nm, dichroic: 590nm, emission filter: 600-650nm). During brightfield imaging a long-pass (600nm) filter was used. The Nikon Perfect Focus System (PFS) was used to prevent loss of focus.

For imaging in unperturbed conditions cells were placed under a prewarmed agar pad (minimal medium, 2% glucose, 1% agarose) and imaged for at least 8 consecutive hours. For each experiment multiple non-overlapping XY positions were recorded and for each position brightfield and fluorescent images were recorded. For imaging of Rpl13a-sfGFP and Rpl26a-sfGFP, 3 brightfield z-axis planes images with a 0.5 µm step were recorded for each XY position every 3 min; one GFP and one RFP fluorescence image were recorded for each position, every 3 min, at the focal plane corresponding to the intermediate brightfield image. For all the other experiments, one brightfield, GFP and RFP image were recorded for every XY position every 5 min.

For experiments involving chemical perturbations using rapamycin, DMSO, MSX, CHX and 1-NM-PP1, exponentially growing cells at OD = 0.1 were incubated for 30 min at 30°C in plastic well plates for inverted fluorescence microscopy (Ibidi) treated with Concanavalin A (1mg/ml). The wells were then washed twice with prewarmed minimal medium (+2% glucose) and placed under the microscope. Imaging settings were as reported above. For each well multiple non-overlapping XY positions were recorded and for each position one brightfield, GFP and RFP image were recorded every 5 min. After at least one hour from the beginning of the imaging, chemicals were added to the wells at the final concentration of 200ng/ml for rapamycin (diluted in DMSO), 2mM for MSX (diluted in H2O), 25 µg/ml for CHX (diluted in H2O) and 500nM for 1-NM-PP1 (diluted in DMSO).

### Image analysis

For each experiment the fluorescence channel images were background-corrected using the rolling ball background subtraction plugin in ImageJ. Cell segmentation and tracking was performed using the semi-automatic ImageJ plugin BudJ (Ferrezuelo et al., 2012). For the calculation of the single-cell N/C ratio of Sfp1-pHtdGFP and Tod6-pHtdGFP, the background-corrected images were subsequently analyzed using a custom-made Python script and the segmentation boundaries detected by BudJ (see below). Mother cells budding events were annotated based on the appearance of a dark spot on the cell membrane, and karyokinesis was annotated based on the first frame when the nucleus of the mother cell and the nucleus of the bud are completely detached. For the calculation of the single-cell synthesis rate of Rpl13a-sfGFP and Rpl26a-sfGFP, the intermediate z-stack brightfield image was used for segmentation and tracking of mother cells, while the brightfield z-stack where the bud was best defined was chosen for the segmentation and tracking of each bud. For the calculation of the synthesis rate, the total (mother + bud) cell volume and the average GFP fluorescence intensity from BudJ were used for subsequent analysis (see below). For all analyses of cell cycle dynamics we considered only mother cells, i.e. cells that had already produced at least one bud.

### Measurement of Sfp1-GFP and Tod6-GFP nuclear-to-cytosolic ratio in single cells

A schematic of the nuclear-to-cytosolic ratio measurement can be found in Supp.Fig.2. For each XY position the segmentation data from BudJ and the corresponding background-corrected images were read in a custom-made Python script. The segmentation information from BudJ was used to generate a mask of the corresponding cell at each time frame. The nucleus of the cell was segmented by applying a simple intensity threshold in the RFP channel within the cell mask. A small and big nuclear mask were calculated as circles of radii 0.48 µm and 1.44 µm respectively, centered at the centroid of the segmented nucleus. Mean GFP intensity in the nucleus was measured by applying the small nuclear mask to the GFP channel and measuring the average pixel intensity inside the mask. Mean cytosolic GFP intensity was calculated by subtracting the big nuclear mask from the cell mask and applying the resulting mask to the GFP channel and calculating the average pixel intensity inside the mask. To quantify the average Sfp1/Tod6/Stb3 N/C ratio in different strains (Fig.3B, Fig.4A,B, Fig.5E,F, Supp.Fig.1A) we quantified the average N/C ratio of mother cells growing under agar pads between minute 200 and minute 400 from the start of imaging.

To compare and average Sfp1 and Tod6 N/C ratio during cell cycles of different length we first split every cell cycle time-series in two parts, one from karyokinesis to the following budding event (G1) and the other from the budding event to the next karyokinesis event (S/G2/M). We then linearly interpolated the first part of every cell cycle time series with a fixed number of equidistant points, and did the same with the second part. The number of points chosen for the interpolation of the first and the second part of each cell cycle time-series were calculated based on the ratio of the average durations of the first part and the second part of the cell cycle. In total, 80 interpolation points were used for each cell cycle. As an example: for an average cell cycle duration of 100min and average G1 and S/G2/M durations of 40min and 60min respectively, the first part of every cell cycle would be interpolated with 32 points and the second with 48 points. For subsequent plotting, we either aligned or averaged the interpolated points of each cell cycle on a common (relative) time scale with 80 steps ranging from 0 to 1.

### Measurement of volume dynamics during the cell cycle, cell cycle phases duration and volume at cell cycle events

To measure the single-cell volume dynamics of WT, ΔPib2 and ΔBcy1 cells during the cell cycle, we used the total volume (mother + bud) time series provided by BudJ, for each cell cycle. We then fitted the volume time series of each cell cycle with a smoothing spline in Matlab to reject artifacts and outliers produced by the segmentation step. To better fit the volume time series at the beginning and at the end of each cell cycle, each volume time series extended with measurements two frames (10min) before the first karyokinesis event and two frames after the next karyokinesis event. Those additional time points were then removed from any subsequent analysis. To compute the volume increase rate during the cell cycle, we differentiated each smoothed volume time series. Averaging of volume increase rate time-series was done in the same way as described above.

G1 duration was defined as the time between karyokinesis and the subsequent budding event. S/G2/M duration we defined as the time taken between budding and the subsequent karyokinesis event. To determine cell volume at budding, we measured with BudJ the volume of mother cells at the first frame where budding was detectable. To measure bud volume at karyokinesis, we measured with BudJ the volumes of buds at the frame where their nucleus completely detached from the nucleus of the mother cell.

### Measurement of Rpl13a and Rpl26a production rate during the cell cycle

To compute the single-cell synthesis rate of Rpl13a-GFP and Rpl26a-GFP during the cell cycle, we first multiplied the total volume (mother + bud) with the mean GFP time-series of the mother cell given by BudJ for each cell cycle, obtaining a GFP abundance time-series for each cell cycle. We assumed that during bud growth, Rpl13a-GFP and Rpl26a-GFP concentration is the same in the mother cell and in the bud, and therefore calculated the mean GFP time series based on the segmentation of the mother cell. Each GFP abundance time series was then fitted with a smoothing spline in Matlab to reject artifacts and outliers produced by the segmentation step. To better fit the GFP abundance time series at the beginning and at the end of each cell cycle, each volume and mean GFP time series were extended with measurements from three frames (9min) before the first karyokinesis and three frames after the second karyokinesis. These extra time points were then removed from any further analysis.

To obtain the maturation-corrected synthesis rate of Rpl13a-GFP and Rpl26a-GFP during the cell cycle, we considered a simple model of protein production and maturation. The immature (i.e. non-fluorescent) protein is synthesized at a (time-varying) rate K_p_(t) and matures in its fluorescent form with a constant rate K_m_. Defining P_i_ as the abundance of the immature protein and P_m_ as the abundance of the mature protein, we obtain the following equations that describe the accumulation of our fluorescent protein:

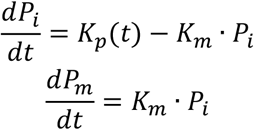

In microscopy experiments, we can actually measure the abundance of the mature protein, i.e. P_m_(t). To estimate the production rate K_p_(t) based on P_m_(t) we use the above equations to arrive at the following:

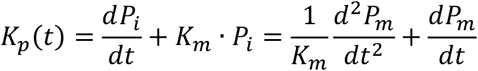

The first derivative of P_m_ was estimated from the smoothing spline of the GFP abundance time series. The first derivative time series was then fitted with a second, closely-fitting smoothing spline to estimate the second derivative of P_m_. In the calculation of K_p_(t) we considered a maturation half time of 6 min for sfGFP (Khmelinskii et al., 2012), leading to K_m_ = 0.116 min^-1^. We did not consider any degradation of RPs during the cell cycle, as there is no evidence for this type of RP regulation. Plotting and averaging of the Rpl13-GFP and Rpl26-GFP synthesis rates during the cell cycle was done in the same way with the Sfp1 and Tod6 time-series. In this case, we fixed the total number of interpolation points at 60.

## Acknowledgements

This work was partly financed by the Netherlands Organization for Scientific Research (NWO) through a VIDI grant to A.M.-A. (project number 016.189.116). We would like to thank Robbie Loewith and David Shore (University of Geneva, Switzerland) for providing strains and plasmids.

## Author Contributions

Conceptualization, methodology and writing, A.M.-A. and P.G.; Software, formal analysis, data curation, visualization, P.G; Investigation, P.G. and M.B.; Resources, P.G., L.-A.V. and M.B.; Funding acquisition, supervision and project administration, A.M.-A.

## Declaration of Interests

The authors declare no competing interests

## Resource Availability

### Lead contact

Further information and requests for resources and reagents should be directed to and will be fulfilled by the Lead Contact, Andreas Milias-Argeitis

### Materials availability

All materials generated in this study are available from the lead contact.

### Data and code availability

Data generated from cell segmentation and tracking, as well as processed microscopy data for all paper figures are available from Mendeley Data:

https://data.mendeley.com/datasets/hvxztrr2m9/draft?a=91da7a1f-88ba-4b17-b13b-c4d9ce5809e0

Python and Matlab^®^ scripts for N/C ratio quantification, cell cycle trace alignment and protein synthesis rate quantification are available at Github: https://github.com/amiliasargeitis/microscopy_scripts

## Notes

### Competing Interest Statement

The authors have declared no competing interest.

