## Supplementary Material for "TORC1 and PKA activity towards ribosome biogenesis oscillates in synchrony with the budding yeast cell cycle"

### Supplementary Materials

**Table S1.** List of yeast strains used in this study. List of target sequences and repairing fragments used for Crispr-Cas9 cloning.

| Yeast strains | | |
|---|---|---|
| YSBN6 | Steve Oliver lab,<br>Cambridge | YSBN6 *wild type* |
| YSBN6<br>Hta2-mRFP | This study | YSBN6<br>*HTA2-mRFP1-Ble* |
| YSBN6<br>Sfp1-pHtdGFP | This study | YSBN6<br>*HTA2-mRFP1-Ble*<br>*SFP1-pHtdGFP-NatMX* |
| YSBN6<br>Tod6-pHtdGFP | This study | YSBN6<br>*HTA2-mRFP1-Ble*<br>*TOD6-pHtdGFP-NatMX* |
| YSBN6<br>Sfp1-mNeonGreen | This study | YSBN6<br>*HTA2-mRFP1-Ble*<br>*SFP1-mNeonGreen-NatMX* |
| YSBN6<br>Tod6-mNeonGreen | This study | YSBN6<br>*HTA2-mRFP1-Ble*<br>*SFP1-mNeonGreen-NatMX* |
| YSBN6<br>Sfp1-pHtdGFP<br>Tpk1-3as | This study | YSBN6<br>*HTA2-mRFP1-Ble*<br>*SFP1-pHtdGFP-NatMX*<br>*TPK1:tpk1_M164G*<br>*TPK2:tpk2_M147G*<br>*TPK3:tpk3_M165G* |
| YSBN6<br>Tod6-pHtdGFP<br>Tpk1-3as | This study | YSBN6<br>*HTA2::mRFP1-Ble*<br>*TOD6::pHtdGFP-NatMX*<br>*TPK1:tpk1_M164G*<br>*TPK2:tpk2_M147G*<br>*TPK3:tpk3_M165G* |
| YSBN6<br>Sfp1-pHtdGFP<br>Sch9$^{2D3E}$ | This study | YSBN6<br>*HTA2-mRFP1-Ble*<br>*SFP1-pHtdGFP-NatMX*<br>*SCH9:sch9*$^{2D3E}$*-KanMX* |
| YSBN6<br>Tod6-pHtdGFP<br>Sch9$^{2D3E}$ | This study | YSBN6<br>*HTA2-mRFP1-Ble*<br>*TOD6-pHtdGFP-NatMX*<br>*SCH9:sch9*$^{2D3E}$*-KanMX* |
| YSBN6<br>Tod6$^{6A}$-pHtdGFP | This study | YSBN6<br>*HTA2-mRFP1-Ble*<br>*HO:TOD6p-tod6*$^{6A}$*-pHtdGFP-KanMX* |
| YSBN6<br>Tod6$^{6A}$-pHtdGFP<br>Tpk1-3as | This study | YSBN6<br>*HTA2-mRFP1-Ble*<br>*HO:TOD6p-tod6*$^{6A}$*-pHtdGFP-KanMX*<br>*TPK1:tpk1_M164G*<br>*TPK2:tpk2_M147G*<br>*TPK3:tpk3_M165G* |

| | | |
|---|---|---|
| YSBN6<br>Sfp1-1-pHtdGFP | This study | YSBN6<br>*HTA2-mRFP1-Ble*<br>*HO:SFP1p-sfp1-1-pHtdGFP-KanMX* |
| YSBN6<br>Sfp1-1-pHtdGFP<br>Tpk1-3as | This study | YSBN6<br>*HTA2-mRFP1-Ble*<br>*HO:SFP1p-sfp1-1-pHtdGFP-KanMX*<br>*TPK1:tpk1_M164G*<br>*TPK2:tpk2_M147G*<br>*TPK3:tpk3_M165G* |
| YSBN6<br>Sfp1-pHtdGFP<br>ΔGtr1<br>ΔGtr2 | This study | YSBN6<br>*HTA2-mRFP1-Ble*<br>*SFP1-pHtdGFP-NatMX*<br>*gtr1Δ*<br>*gtr2Δ* |
| YSBN6<br>Tod6-pHtdGFP<br>ΔGtr1<br>ΔGtr2 | This study | YSBN6<br>*HTA2-mRFP1-Ble*<br>*TOD6-pHtdGFP-NatMX*<br>*gtr1Δ*<br>*gtr2Δ* |
| YSBN6<br>Sfp1-pHtdGFP<br>ΔPib2 | This study | YSBN6<br>*HTA2-mRFP1-Ble*<br>*SFP1-pHtdGFP-NatMX*<br>*pib2Δ::KanMX* |
| YSBN6<br>Tod6-pHtdGFP<br>ΔPib2 | This study | YSBN6<br>*HTA2-mRFP1-Ble*<br>*TOD6-pHtdGFP-NatMX*<br>*pib2Δ::KanMX* |
| YSBN6<br>Sfp1-pHtdGFP<br>ΔPib2<br>$Gtr1^{Q65L}$<br>$Gtr2^{S23L}$ | This study | YSBN6<br>*HTA2-mRFP1-Ble*<br>*SFP1-pHtdGFP-NatMX*<br>*pib2Δ::KanMX*<br>*GTR1:gtr1_Q65L*<br>*GTR2:gtr2_S23L* |
| YSBN6<br>Tod6-pHtdGFP<br>ΔPib2<br>$Gtr1^{Q65L}$<br>$Gtr2^{S23L}$ | This study | YSBN6<br>*HTA2-mRFP1-Ble*<br>*TOD6-pHtdGFP-NatMX*<br>*pib2Δ::KanMX*<br>*GTR1:gtr1_Q65L*<br>*GTR2:gtr2_S23L* |
| YSBN6<br>Sfp1-pHtdGFP<br>ΔBcy1 | This study | YSBN6<br>*HTA2-mRFP1-Ble*<br>*SFP1-pHtdGFP-NatMX*<br>*bcy1Δ::KanMX* |
| YSBN6<br>Tod6-pHtdGFP<br>ΔBcy1 | This study | YSBN6<br>*HTA2-mRFP1-Ble*<br>*TOD6-pHtdGFP-NatMX*<br>*bcy1Δ::KanMX* |
| YSBN6<br>Sfp1-pHtdGFP<br>$Ras2^{A18V19}$<br>$Gpa2^{A273}$ | This study | YSBN6<br>*HTA2-mRFP1-Ble*<br>*SFP1-pHtdGFP-NatMX*<br>*RAS2:ras2_A18V19*<br>*GPA2:gpa2_A273* |

| | | |
|---|---|---|
| YSBN6<br>Tod6-pHtdGFP<br>Ras2$^{A18V19}$<br>Gpa2$^{A273}$ | This study | YSBN6<br>*HTA2-mRFP1-Ble*<br>*TOD6-pHtdGFP-NatMX*<br>*RAS2:ras2_A18V19*<br>*GPA2:gpa2_A273* |
| YSBN6<br>Rpl13a-sfGFP | This study | YSBN6<br>*HTA2.mRFP1-Ble*<br>*RPL13A.sfGFP-KanMX* |
| YSBN6<br>Rpl26a-sfGFP | This study | YSBN6<br>*HTA2-mRFP1-Ble*<br>*RPL26A-sfGFP-KanMX* |
| CRISPR-Cas9 target sequences and repairing fragments | | |
| Cas9 target sequence for Tpk1:<br>TGAAAAATTGCTGAGCATCT | This study | |
| Cas9 target sequence for Tpk2:<br>GTGATGGATTATATCGAAGG | This study | |
| Cas9 target sequence for Tpk3:<br>GTAATGGCCTACATTGAAGG | This study | |
| Cas9 target sequence for Gtr1:<br>GAATATGACTCTAAATCTGT | This study | |
| Cas9 target sequence for Gtr2:<br>TGGTTTTGTTGATGGGCGTA | This study | |
| Cas9 target sequence for Ras2:<br>TACAAGCTAGTCGTCGTTGG | This study | |
| Cas9 target sequence for Gpa2:<br>CTTAATATGACCTGCTGGGT | This study | |
| Gtr1KO repairing fragment:<br>AGGTATCTTACACAGGAGTGAAGGCCA<br>TCAAAATCACGTTTATCAATCGACAATTT<br>AGTACTGAGGTGAGTAGACGAAACATTC<br>GGCAATTGAGTGTTTGCGGGGCATAAG<br>AATTATAAA | This study | |
| Gtr2KO repairing fragment:<br>ACCGATTAACATCCACAGATTAACAAAA<br>CTCCAGGACAACGGTACTAATACACATA<br>CAACAAGACGTAAGGCATGAAAATATTA<br>GGGTATATAGATACATATTGAAAATGAT<br>AGTAGAGC | This study | |
| Gtr1_Q65L repairing fragment:<br>GCCACCATTGATGTAGAGCACTCCCATT<br>TGAGATTTCTTGGGAATATGACTCTTAA<br>CCTCTGGGACTGTGGTGGGCTGGACGT<br>GTTTATGGAGAATTATTTCACCAAGCAA<br>AAAGACCAC | This study | |
| Gtr2_S23L repairing fragment:<br>CCAGGACAACGGTACTAATACACATACA<br>ACATGAGTTTAGAGGCTACAGATTCCAA<br>GGCAATGGTTTTGTTGATGGGCGTAAG<br>AAGATGTGGAAAATTATCCATTTGTAAA<br>GTTGTTTTT | This study | |

| | |
|---|---|
| Ras2_A18V19 repairing fragment:<br>GAATTGAAAGGAGATATACAGAAAAAAA<br>AATGCCTTTGAACAAGTCGAACATAAGA<br>GAGTACAAGCTAGTTGTTGTCGGAGCT<br>GTTGGTGTTGGTAAATCTGCTTTGACCA<br>TACAATTGACCCAATCGCACTTTGTAGA<br>TGAATACGATCCCACAATTGA | This study |
| Gpa2_A273 repairing fragment:<br>CGAAGTTCTATCTAATGGACTCGACTCC<br>TTACTTCATGGAAAATTTCACCAGGATC<br>ACTTCGCCCAATTACAGACCCACTCAAC<br>AAGACATATTAAGATCGGCTCAGATGAC<br>GTCAGGGATTTTTGACACCGTCATTGAT<br>ATGGGGTCGGATATCAAGATGCATATTT<br>ACGACGTGGGTG | This study |
| Tpk1_M164G repairing fragment:<br>TTTCTATCGTAACACATCCGTTTATTATT<br>AGAATGTGGGGGACTTTCCAAGATGCT<br>CAGCAAATTTTCATGATTGGTGATTATAT<br>TGAAGGTGGAGAATTGTTTTCTTTGTTA<br>AGGAAATCCCAAAGATTTCCCAATCCTG<br>TCGCTAAATTTTACGCAGCGGAAGTTTG<br>TTTAGCTTTGGAGTACTTGCATAGCAAG<br>GACATTATTTATAGGGATTTGAA | This study |
| Tpk2_M147G repairing fragment:<br>TGATTAGAATGTGGGGTACGTTTCAAGA<br>TGCTAGGAATATCTTTATGGTGGGTGAT<br>TATATAGAGGGTGGTGAACTTTTCTCGT<br>TACTGAGAAAGTCACAAAGATTTCCTAA<br>TCCTGTAG | This study |
| Tpk3_M165G repairing fragment:<br>CATCATTCGAATGTGGGGAACGTTCCAA<br>GATTCTCAGCAAGTTTTCATGGTAGGTG<br>ACTACATCGAG<br>GGTGGTGAATTATTTTCTTTACTACGTAA<br>ATCTCAAAGATTTCCCAACCCAGT | This study |

### Supplementary figures

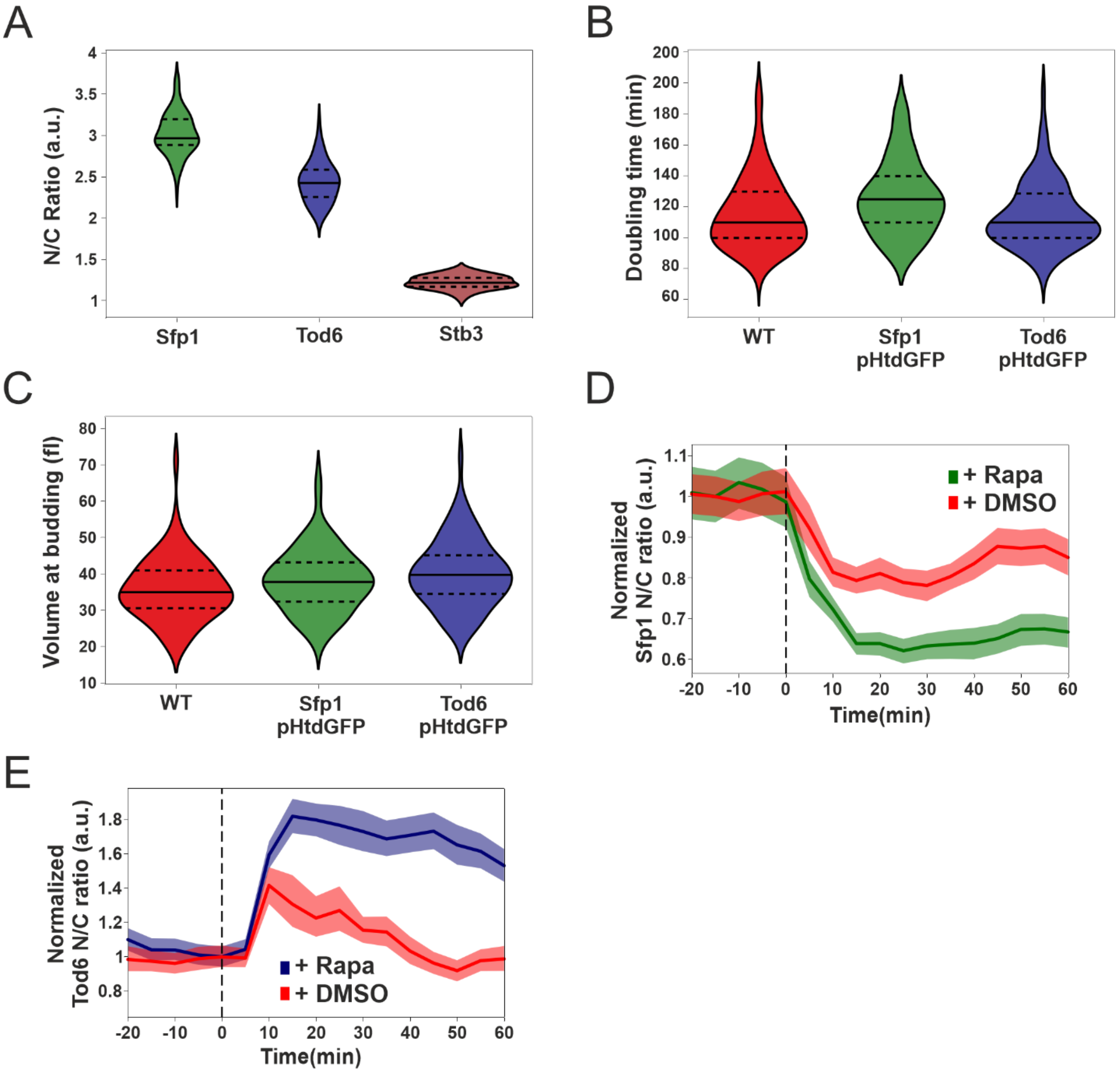

**Figure S1**. Related to Figure 1

**A.** Distributions of single-cell N/C ratios for Sfp1-pHtdGFP (n=70), Tod6-pHtdGFP (n=64) and Stb3-pHtdGFP (n=53). Median (continuous line) and 25th and 75th percentiles (dashed lines) are also displayed. **B.** Doubling time (karyokinesis-to-karyokinesis) distributions for single mother cells of the wild type (n = 100), Sfp1-pHtdGFP (n = 72) and Tod6-pHtdGFP (n = 94) backgrounds. Median (continuous line) and 25th and 75th percentiles (dashed lines) are also displayed. Statistical comparison, two-tailed Mann-Whitney test: Sfp1-WT p-value = 0.002, effect size (rank-biserial correlation) r = 0.26, Tod6-WT p-value = 0.9, r = 0.004 **C.** Distributions of single-cell volumes at budding for mother cells of the wild type (n = 68), Sfp1-pHtdGFP (n =

71) and Tod6-pHtdGFP (n = 77) backgrounds. Median (continuous line) and 25th and 75th percentiles (dashed lines) are also displayed. Statistical comparison, Mann-Whitney test: Sfp1-WT p-value = 0.088, r = 0.16, Tod6-WT p-value = 0.004, r = 0.27. **D.** Normalized N/C ratio dynamics of Sfp1 in response to rapamycin (n=56) and its vehicle (DMSO)(n=69). Cells were attached to plastic wells treated with ConcanavalinA (1mg/ml) and imaged every 5 min. Rapamycin (200ng/ml final) and DMSO (0.6% v/v) were added at t = 0. N/C ratio traces were normalized at their value at t=0 to facilitate comparisons. The bands denote the 95% confidence interval for the mean. **E.** Normalized N/C ratio dynamics of Tod6 in response to rapamycin (n=61) and its vehicle (DMSO) (n=59). Cells were attached to plastic wells treated with ConcanavalinA (1mg/ml) and imaged every 5 min. Rapamycin (200ng/ml final) and DMSO (0.6% v/v) were added at t = 0. N/C ratio traces were normalized at their value at t=0 to facilitate comparisons. The bands denote the 95% confidence interval for the mean.

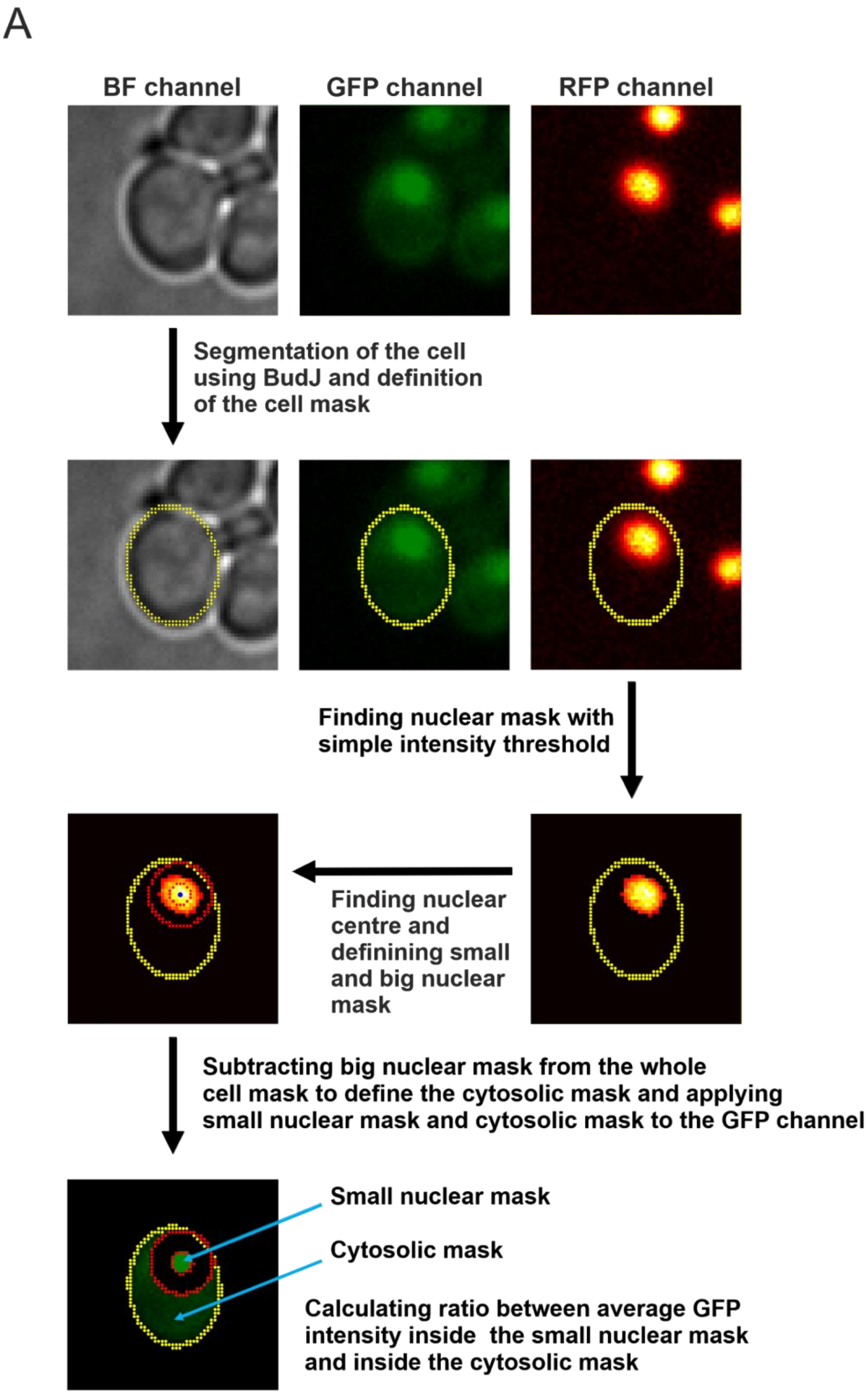

**Figure S2.** Related to Figure 1,2,3,4,5,
Schematic representation of the pipeline to calculate Sfp1 and Tod6 N/C ratio in single cells

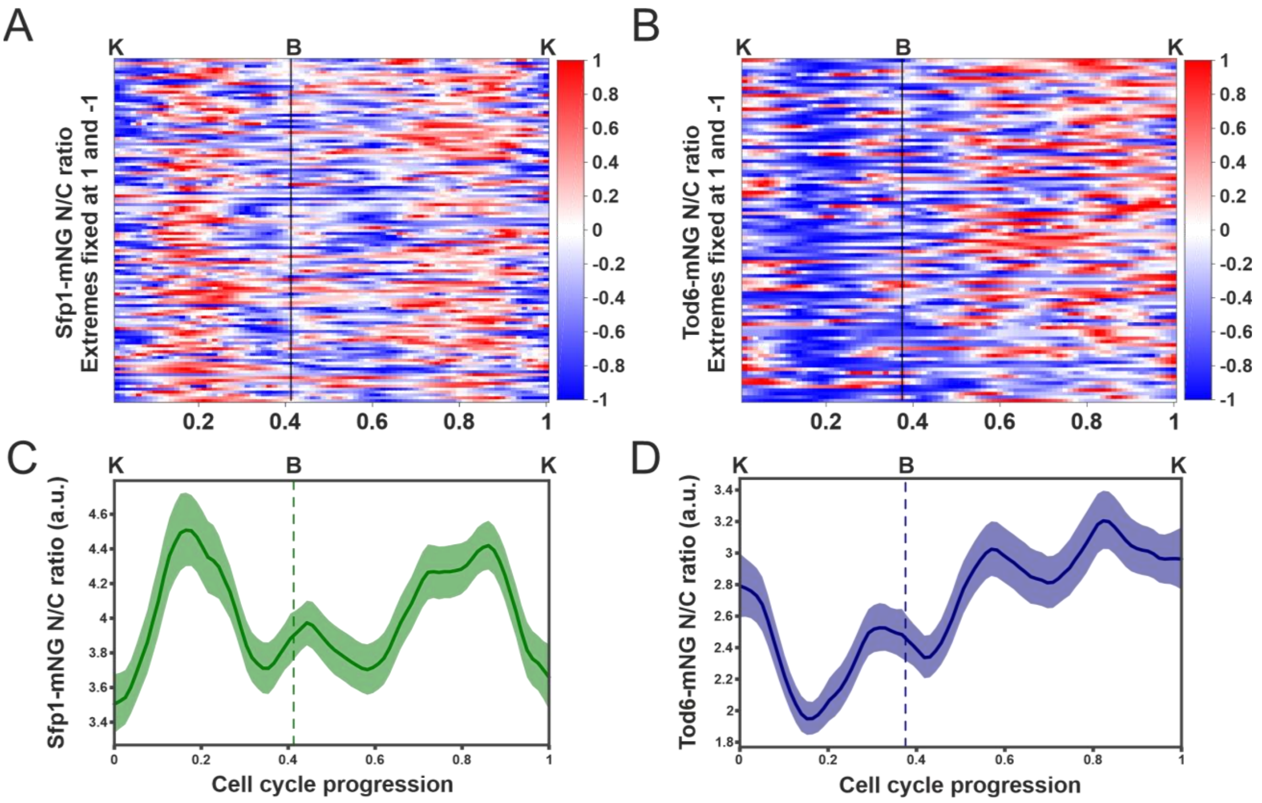

**Figure S3**. Related to Figure 2

**A.** Heatmap of Sfp1-mNeonGreen N/C ratio in individual cell cycles (n = 119). The cell cycles traces were interpolated and aligned as described in Fig.2A. For each cell cycle, the Sfp1 N/C ratio was normalized by assigning its maximum to 1 and its minimum to -1, to facilitate the identification of peaks and troughs. **B**. Heatmap of Tod6-mNeonGreen N/C ratio in individual cell cycles (n = 99). The cell cycles traces were interpolated and aligned as described in Fig.2A. For each cell cycle, the Tod6 N/C ratio was normalized by assigning its maximum to 1 and its minimum to -1, to facilitate the identification of peaks and troughs. **C.** Average Sfp1 N/C ratio dynamics for the cells shown in **A**. The averages were calculated without normalization of the single-cell data. The bands denote the 95% confidence interval for the mean. **D**. Average Tod6 N/C ratio dynamics for the cells shown in **B**. The averages were calculated without normalization of the single-cell data. The bands denote the 95% confidence interval for the mean.

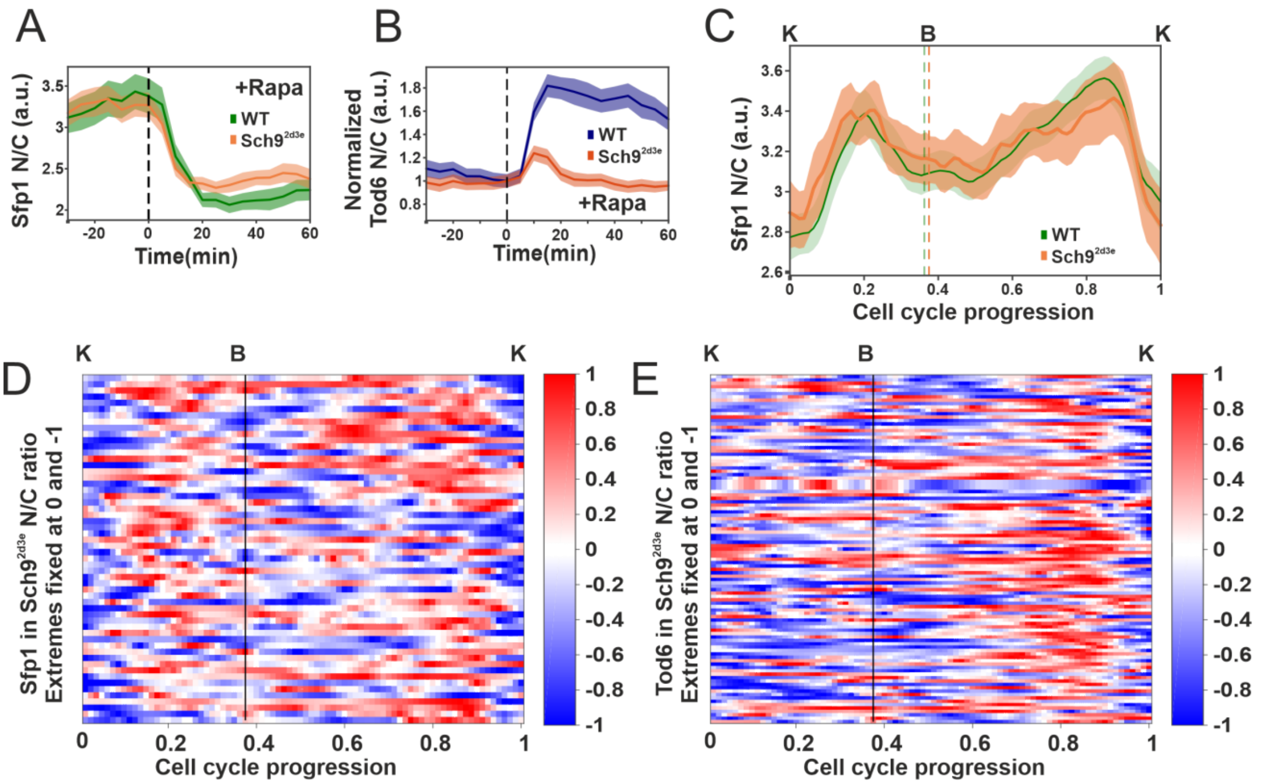

**Figure S4**. Related to Figure 3

**A.** N/C ratio dynamics of Sfp1 in WT(n=56) and in Sch9_2d3e (n=68) cells in response to rapamycin. Rapamycin (200ng/ml final) was added at time t=0. The bands denote the 95% confidence interval for the mean. **B.** Normalized N/C ratio dynamics of Tod6 in WT (n=61) and in Sch9_2d3e (n=76) cells in response to rapamycin. Rapamycin (200ng/ml final) was added at time t=0. N/C ratio single cell traces were normalized to their value at t=0 to facilitate comparison. The bands denote the 95% confidence interval for the mean. **C.** Average Sfp1 N/C ratio dynamics in WT (n=149) and in Sch9_2d3e cells (n = 56 cells). The averages were calculated without normalization of the single-cell data. Individual cell cycles traces were interpolated and aligned as described in Fig.2A. Bands denote the 95% confidence interval for the mean. **D**. Heatmap of Sfp1 in Sch9_2d3e cells N/C ratio in individual cell cycles (n = 56). The cell cycles traces were interpolated and aligned as described in Fig.2A. For each cell cycle, the Sfp1 N/C ratio was normalized by assigning its maximum to 1 and its minimum to -1, to facilitate the identification of peaks and troughs. **E.** Heatmap of Tod6 in Sch9_2d3e N/C ratio in individual cell cycles (n = 103). The cell cycles traces were interpolated and aligned as described in Fig.2A. For each cell cycle, the Tod6 N/C ratio was normalized by assigning its maximum to 1 and its minimum to -1, to facilitate the identification of peaks and troughs.

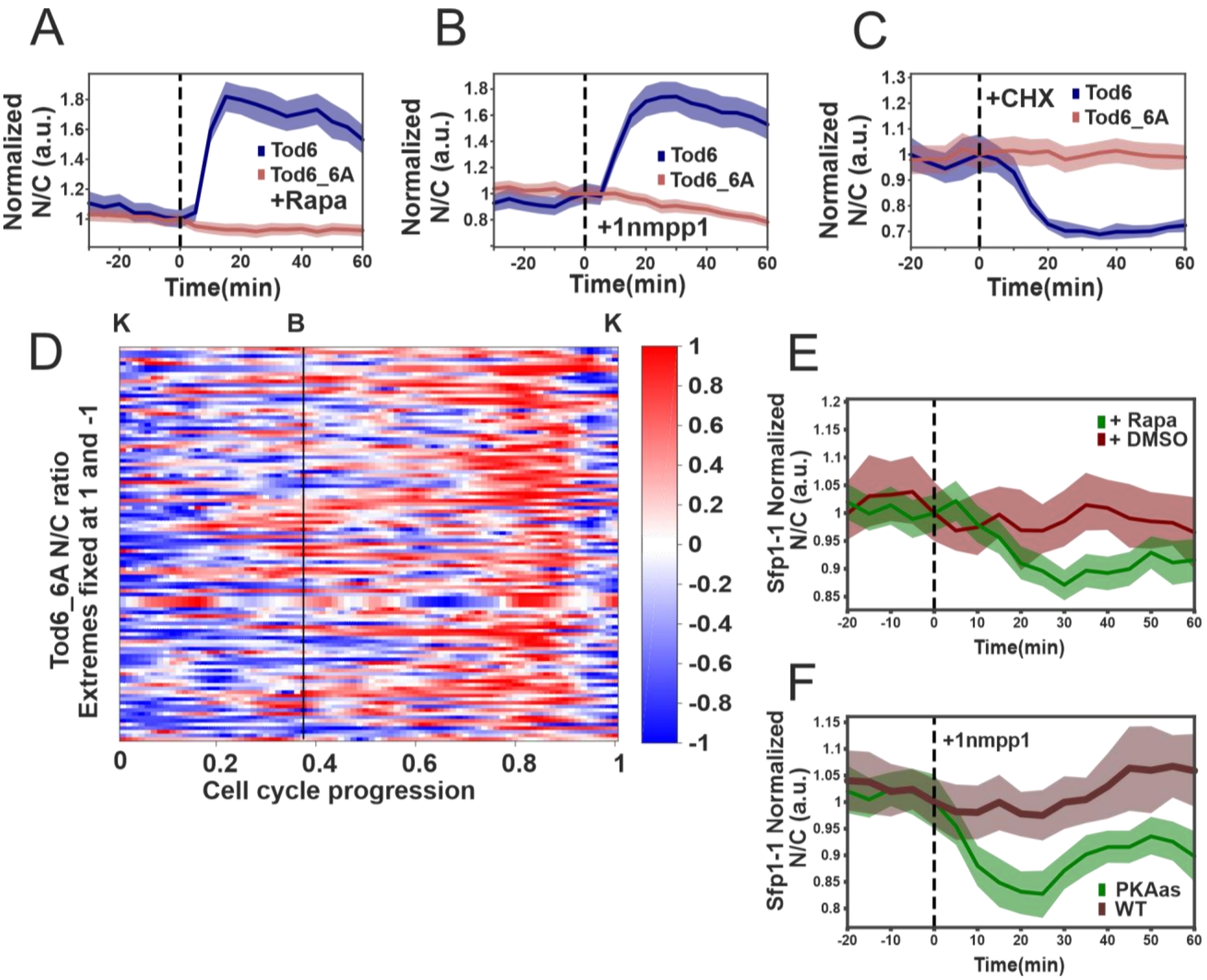

**Figure S5**. Related to Figure 3

**A.** N/C ratio changes of Tod6(n= 61) and Tod6_6A(n=120) in response to rapamycin. Rapamycin (200ng/ml final) was added at time t=0. N/C ratio single cell traces were normalized at their value at t=0 to facilitate comparison. The bands denote the 95% confidence interval for the mean. **B.** N/C ratio changes of Tod6(n=63) and Tod6_6A(n=75) in response to addition of 1-NM-PP1 in PKAas cells. 1-NM-PP1 (500nM final) was added at time t=0. N/C ratio single cell traces were normalized at their value at t=0 to facilitate comparison. The bands denote the 95% confidence interval for the mean. **C.** N/C ratio changes of Tod6(n=56) and Tod6_6A(n=65) in response to addition of CHX. CHX (25ug/ml final) was added at time t=0. N/C ratio single cell traces were normalized at their value at t=0 to facilitate comparison. The bands denote the 95% confidence interval for the mean. **D.** Heatmap of Tod6_6A N/C ratio in individual cell cycles (n = 109). Single-cell traces of Tod6_6A localization were aligned and interpolated as described in Fig. 2A. For each cell cycle, the Tod6_6A N/C ratio was normalized by assigning its maximum to 1 and its minimum to -1, to facilitate the identification of peaks and troughs. **E.** N/C ratio changes of Sfp1-1 in response to rapamycin (n=71) and its vehicle DMSO (n=19). Rapamycin (200ng/ml final) and DMSO (0.66% v/v) were at time t=0. N/C ratio single cell traces were normalized at their value at t=0. The bands denote the 95% confidence interval for the mean. **F.**

N/C ratio changes of Sfp1-1 in response to addition of 1-NM-PP1 in PKAas cells (n=24) and WT cells (n=22). 1-NM-PP1 (500nM final) was added at time t=0. N/C ratio single cell traces were normalized at their value at t=0. The bands denote the 95% confidence interval for the mean.

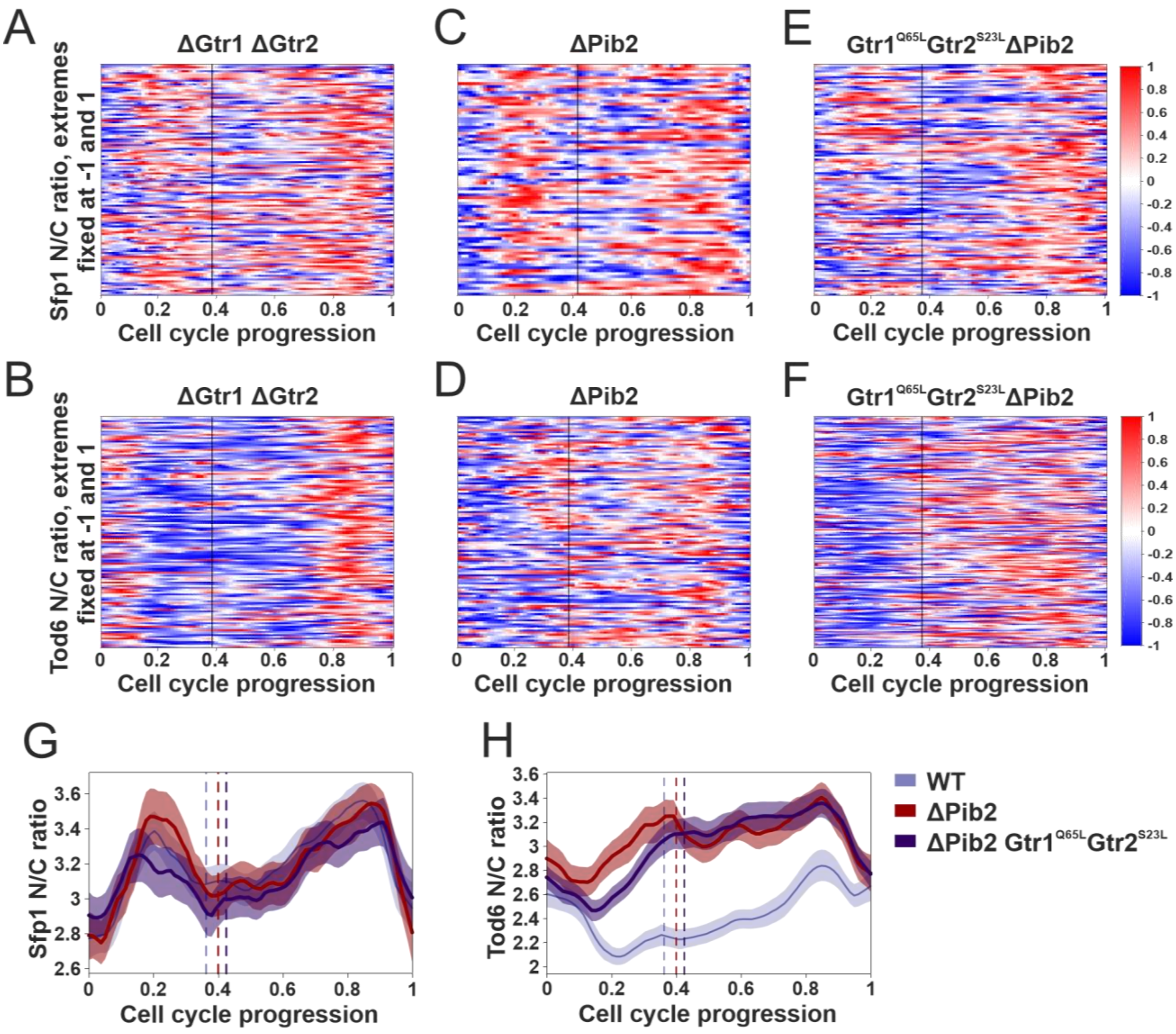

**Figure S6**. Related to Figure 4

**A.** Heatmap of Sfp1 N/C ratio in individual cell cycles (n = 122) in ΔGtr1ΔGtr2 cells. Single-cell traces of Sfp1 localization were aligned and interpolated as described in Fig. 2A. For each cell cycle, the Sfp1 N/C ratio was normalized by assigning its maximum to 1 and its minimum to -1. **B.** Heatmap of Tod6 N/C ratio in individual cell cycles (n = 136) in ΔGtr1ΔGtr2 cells. Single-cell traces of Tod6 localization were aligned and interpolated as described in Fig. 2A. For each cell cycle, the Sfp1 N/C ratio was normalized by assigning its maximum to 1 and its minimum to -1. **C.** Heatmap of Sfp1 N/C ratio in individual cell cycles (n = 79) in ΔPib2 cells. Single-cell traces of Sfp1 localization were aligned and interpolated as described in Fig. 2A. For each cell cycle, the Sfp1 N/C ratio was normalized by assigning its maximum to 1 and its minimum to -1. **D.** Heatmap of Tod6 N/C ratio in individual cell cycles (n = 113) in ΔPib2 cells. Single-cell traces of Tod6 localization were aligned and interpolated as described in Fig. 2A. For each cell cycle, the Sfp1 N/C ratio was normalized by assigning its maximum to 1 and its minimum to -1. **E.** Heatmap of Sfp1 N/C ratio in individual cell cycles (n = 118) in Gtr1_Q65L_Gtr2_S23L_ΔPib2 cells. Single-cell traces of Sfp1 localization were aligned and interpolated as described in Fig.

2A. For each cell cycle, the Sfp1 N/C ratio was normalized by assigning its maximum to 1 and its minimum to -1. **F.** Heatmap of Tod6 N/C ratio in individual cell cycles (n = 181) in Gtr1_Q65L_Gtr2_S23L_ΔPib2 cells. Single-cell traces of Tod6 localization were aligned and interpolated as described in Fig. 2A. For each cell cycle, the Sfp1 N/C ratio was normalized by assigning its maximum to 1 and its minimum to -1. **G.** Average Sfp1 N/C ratio dynamics in WT (n=149), ΔPib2 (n=79) and Gtr1_Q65L_Gtr2_S23L_ΔPib2 (n=118) cells. The averages were calculated without normalization of the single-cell data. The cell cycles were interpolated and aligned as described in Fig.2A. The bands denote the 95% confidence interval for the mean. **H.** Average Tod6 N/C ratio dynamics in WT (n=161), ΔPib2 (n=136) and Gtr1_Q65L_Gtr2_S23L_ΔPib2 (n=181) cells. The averages were calculated without normalization of the single-cell data. The cell cycles were interpolated and aligned as described in Fig.2A. The bands denote the 95% confidence interval for the mean.

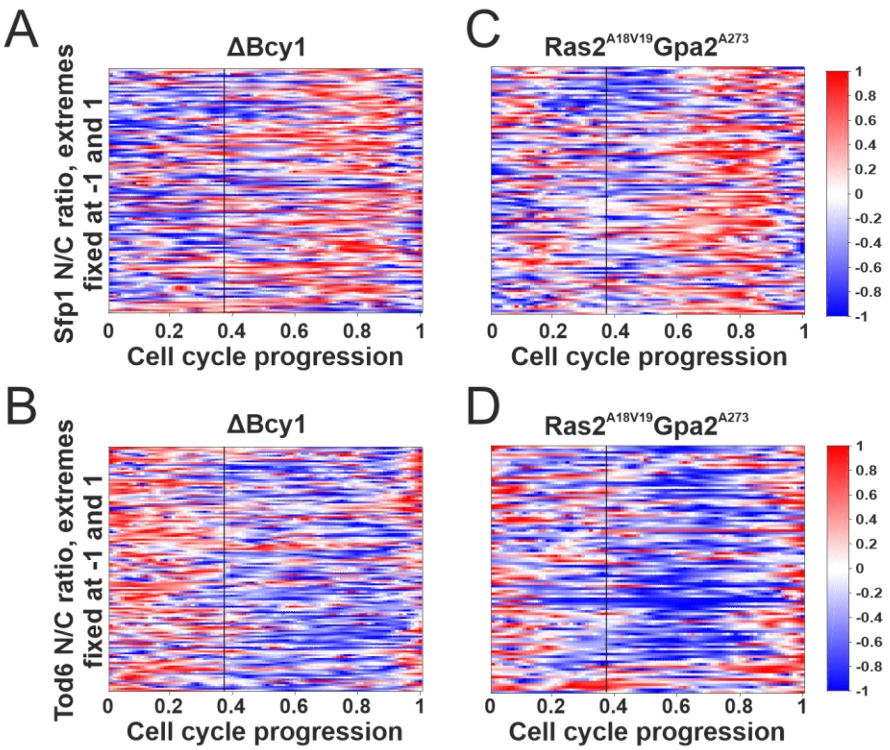

**Figure S7**. Related to Figure 5

**A.** Heatmap of Sfp1 N/C ratio in individual cell cycles (n = 125) in ΔBcy1 cells. Single-cell traces of Sfp1 localization were aligned and interpolated as described in Fig. 2A. For each cell cycle, the Sfp1 N/C ratio was normalized by assigning its maximum to 1 and its minimum to -1. **B.** Heatmap of Tod6 N/C ratio in individual cell cycles (n = 131) in ΔBcy1 cells. Single-cell traces of Tod6 localization were aligned and interpolated as described in Fig. 2A. For each cell cycle, the Sfp1 N/C ratio was normalized by assigning its maximum to 1 and its minimum to -1. **C.** Heatmap of Sfp1 N/C ratio in individual cell cycles (n = 111) in Ras2_A18V19_Gpa2_A273 cells. Single-cell traces of Sfp1 localization were aligned and interpolated as described in Fig. 2A. For each cell cycle, the Sfp1 N/C ratio was normalized by assigning its maximum to 1 and its minimum to -1. **D.** Heatmap of Tod6 N/C ratio in individual cell cycles (n = 100) in Ras2_A18V19_Gpa2_A273 cells. Single-cell traces of Tod6 localization were aligned and interpolated as described in Fig. 2A. For each cell cycle, the Sfp1 N/C ratio was normalized by assigning its maximum to 1 and its minimum to -1.

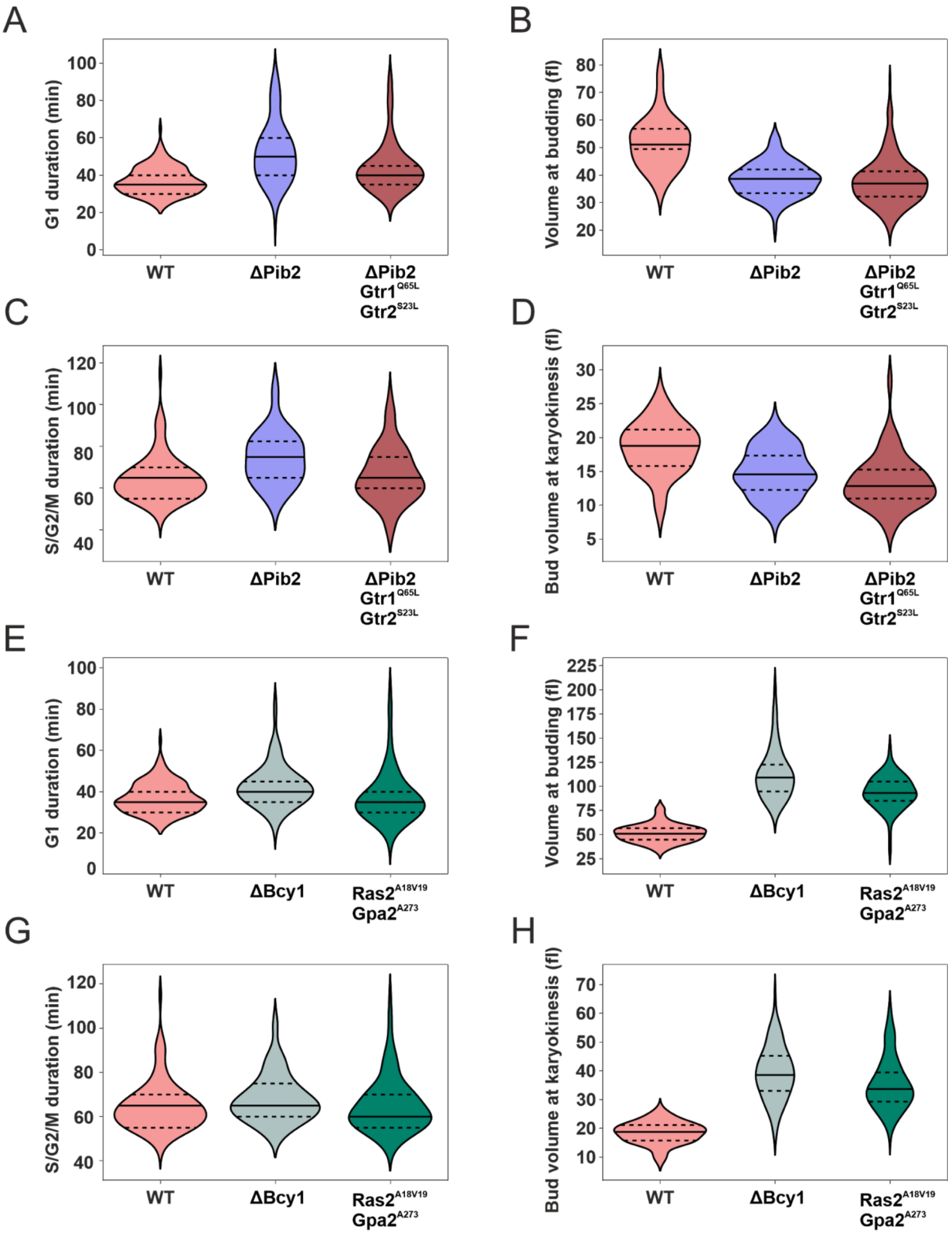

**Figure S8**. Related to Figure 6

**A**. G1 duration distributions of single cells in the wild-type (n = 145), ΔPib2 (n = 112) and ΔPib2_Gtr1$^{Q65L}$Gtr2$^{S23L}$ (n = 183) mutants. Median (continuous line) and 25th and 75th percentiles (dashed lines) are also displayed. G1 was defined as the interval between karyokinesis and bud appearance. This definition slightly overestimates the actual G1 duration. Statistical comparison, two-tailed Mann-Whitney test: ΔPib2-WT $p = 1.5\cdot10^{-17}$,r = 0.61, ΔPib2_Gtr1$^{Q65L}$Gtr2$^{S23L}$-WT $p = 1.1\cdot10^{-7}$ , r = 0.34. **B**. Distributions of volume at the moment of bud appearance in the wild-type (n = 150) and the ΔPib2 (n = 122) and ΔPib2_Gtr1$^{Q65L}$Gtr2$^{S23L}$ (n = 183) mutants. Median (continuous line) and 25th and 75th percentiles (dashed lines) are also displayed. Statistical comparison, two-tailed Mann-Whitney test: ΔPib2-WT $p = 5.6\cdot10^{-29}$, r = 0.78, ΔPib2_Gtr1$^{Q65L}$Gtr2$^{S23L}$-WT $p = 1.1\cdot10^{-32}$ , r = 0.76. **C**. S/G2/M duration distributions in the wild-type (n = 113) and the ΔPib2 (n = 103) and ΔPib2_Gtr1$^{Q65L}$Gtr2$^{S23L}$ (n = 98) mutants. Median (continuous line) and 25th and 75th percentiles (dashed lines) are also displayed. S/G2/M was defined as the interval between bud appearance and karyokinesis. This definition slightly underestimates the actual duration of these phases. Statistical comparison, two-tailed Mann-Whitney test: ΔPib2-WT $p = 1\cdot10^{-9}$, r = 0.47, ΔPib2_Gtr1$^{Q65L}$Gtr2$^{S23L}$-WT p = 0.26, r = 0.09 **D**. Distributions of bud volume at the moment of karyokinesis in the wild-type (n = 109) and the ΔPib2 (n = 103) and ΔPib2_Gtr1$^{Q65L}$Gtr2$^{S23L}$ (n = 102) mutants. Median (continuous line) and 25th and 75th percentiles (dashed lines) are also displayed. Statistical comparison, two-tailed Mann-Whitney test: ΔPib2-WT $p = 3.2\cdot10^{-11}$, r = 0.53, ΔPib2_Gtr1$^{Q65L}$Gtr2$^{S23L}$-WT $p = 2.6\cdot10^{-17}$ , r = 0.67. **E**. G1 duration distributions of single cells in the wild-type (n = 145), ΔBcy1 (n = 135) and Ras2$^{A18V19}$Gpa2$^{A273}$ (n = 98) mutants. Median (continuous line) and 25th and 75th percentiles (dashed lines) are also displayed. G1 was defined as the interval between karyokinesis and bud appearance. This definition overestimates the actual G1 duration. Statistical comparison, two-tailed Mann-Whitney test: ΔBcy1-WT $p = 4.7\cdot10^{-7}$,r = 0.34, Ras2$^{A18V19}$Gpa2$^{A273}$-WT p = 0.25 , r = 0.08. **F**. Distributions of volume at the moment of bud appearance in the wild-type (n = 150) and the ΔBcy1 (n = 135) and Ras2$^{A18V19}$Gpa2$^{A273}$ (n = 98) mutants. Median (continuous line) and 25th and 75th percentiles (dashed lines) are also displayed. Statistical comparison, two-tailed Mann-Whitney test: ΔBcy1-WT $p = 5.9\cdot10^{-48}$, r = 0.99, Ras2$^{A18V19}$Gpa2$^{A273}$-WT $p = 1.06\cdot10^{-36}$ , r = 0.95. **G**. S/G2/M duration distributions in the wild-type (n = 113) and the ΔBcy1 (n = 113) and Ras2$^{A18V19}$Gpa2$^{A273}$ (n = 125) mutants. Median (continuous line) and 25th and 75th percentiles (dashed lines) are also displayed. S/G2/M was defined as the interval between bud appearance and karyokinesis. This definition slightly underestimates the actual duration of these phases. Statistical comparison, two-tailed Mann-Whitney test: ΔBcy1-WT p =0.004, r = 0.21, Ras2$^{A18V19}$Gpa2$^{A273}$-WT p = 0.93, r = 0.29 **H**. Distributions of bud volume at the moment of karyokinesis in the wild-type (n = 109) and the ΔBcy1 (n = 110) and Ras2$^{A18V19}$Gpa2$^{A273}$ (n = 123) mutants. Median (continuous line) and 25th and 75th percentiles (dashed lines) are also displayed. Statistical comparison, two-tailed Mann-Whitney test: ΔBcy1-WT $p = 1.9\cdot10^{-35}$, r = 0.97, Ras2$^{A18V19}$Gpa2$^{A273}$-WT $p = 1.9\cdot10^{-37}$ , r = 0.97

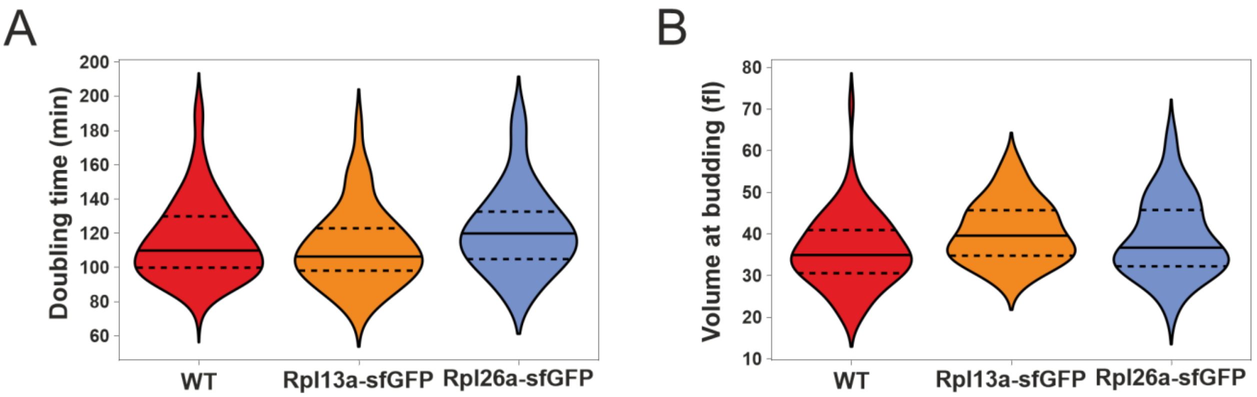

**Figure S9**. Related to Figure 7

**A.** Doubling time (karyokinesis-to-karyokinesis) distributions for single mother cells of the wild type (n = 100), Rpl13a-sfGFP (n = 44) and Rpl26a-sfGFP (n = 48) backgrounds. Median (continuous line) and 25th and 75th percentiles (dashed lines) are also displayed. Statistical comparison, Mann-Whitney test: Rpl13a-WT p-value = 0.36, r = 0.09, Rpl26a-WT p-value = 0.13, r = 0.15 **B.** Distributions of single-cell volumes at budding for mother cells of the wild type (n = 68), Rpl13a-sfGFP (n = 58) and Rpl26a-sfGFP (n = 62) backgrounds. Median (continuous line) and 25th and 75th percentiles (dashed lines) are also displayed. Statistical comparison, Mann-Whitney test: Rpl13a-WT p-value = 0.001, r = 0.33, Rpl26a-WT p-value = 0.07, r = 0.18
